# The fission yeast methyl phosphate capping enzyme Bmc1 guides 2’-O-methylation of the U6 snRNA

**DOI:** 10.1101/2023.01.27.525755

**Authors:** Jennifer Porat, Viktor A. Slat, Stephen D. Rader, Mark A. Bayfield

**Affiliations:** Department of Biology, York University, Toronto, Canada; Department of Biochemistry and Molecular Biology, University of British Columbia, Vancouver, Canada; Department of Chemistry and Biochemistry, University of Northern British Columbia, Prince George, Canada

**Keywords:** RNA, RNA methylation, small nuclear RNA, spliceosome, splicing

## Abstract

Splicing requires the tight coordination of dynamic spliceosomal RNAs and proteins. U6 is the only spliceosomal RNA transcribed by RNA Polymerase III and undergoes an extensive maturation process. In humans and fission yeast, this includes addition of a 5’ γ-monomethyl phosphate cap by members of the Bin3/MePCE family. Previously, we have shown that the Bin3/MePCE homolog Bmc1 is recruited to the *S. pombe* telomerase holoenzyme by the LARP7 family protein Pof8, where it acts in a catalytic-independent manner to protect the telomerase RNA and facilitate holoenzyme assembly. Here, we show that Bmc1 and Pof8 also interact in a U6-containing snRNP. We demonstrate that Bmc1 and Pof8 promote 2’-O-methylation of U6 and identify and characterize a non-canonical snoRNA that guides this methylation. Further, we show that fission yeast strains deleted of Bmc1 or Pof8 show altered U6 snRNP assembly patterns, supporting a more general role for these factors in guiding noncoding RNP assembly beyond the telomerase RNP. These results are thus consistent with a novel role for Bmc1/MePCE family members in stimulating U6 post-transcriptional modifications.

## Introduction

Pre-mRNA splicing, comprised of intron excision and subsequent exon ligation, relies on dynamic RNA-RNA and RNA-protein interactions in the spliceosome (reviewed in (Wilkinson, Charenton and Nagai, 2020)). The spliceosome contains upwards of 100 proteins (Cvitkovic and Jurica, 2013) and 5 uridylate-rich small nuclear RNAs (snRNAs): U1, U2, U4, U5, and U6. The U6 snRNA, which forms part of the catalytic core of the spliceosome (Fica *et al*., 2013), undergoes several conformational changes during pre-spliceosome assembly and splicing catalysis, which enables its interaction with other spliceosomal RNAs and the switch between a catalytically active and inactive state (Eysmont *et al*., 2019). As such, U6 biogenesis and maturation is complex and tightly regulated to ensure correct functioning in the spliceosome (reviewed in (Didychuk, Butcher and Brow, 2018)).

In addition to being the most highly conserved of the snRNAs, U6 is also the only snRNA transcribed by RNA Polymerase III (RNAP III) (Brow and Guthrie, 1988). Transcription of U6 by RNAP III is associated with the addition of a 5’ γ-monomethyl phosphate cap catalyzed by enzymes of the Bin3/MePCE (methyl phosphate capping enzyme) family (Singh and Reddy, 1989; Jeronimo *et al*., 2007; Cosgrove *et al*., 2012) and a 3’ uridylate tail. U6 contains a 5’ stem loop critical for 5’ capping (Singh, R., Gupta, S., and Reddy, 1990), as well as an internal stem loop (ISL) that forms during splicing catalysis. The ISL is mutually exclusive with U4/U6 base pairing that occurs in pre-spliceosome snRNPs (Huppler *et al*., 2002). U6 also contains 2’-O-methylated, pseudouridylated, and m6A-modified nucleotides, with pseudouridines largely present towards the 5’ end and 2’-O-methylations tending to cluster in the ISL (Reddy and Busch, 1988; Gu *et al*., 1996). Moreover, U6 maturation in the fission yeast *Schizosaccharomyces pombe* involves the splicing of an mRNA-type intron, thought to arise from reverse splicing, as the intron is located near the catalytic nucleotides of U6 (Brow and Guthrie, 1989; Potashkin and Frendewey, 1989; Tani and Ohshima, 1989, 1991; Frendewey *et al*., 1990). Most information about the timing of U6 processing events has come from elegant studies in budding yeast (reviewed in (Didychuk, Butcher and Brow, 2018)). However, since budding yeast U6 lacks 2’-O-methylations and a Bmc1 homolog (Deragon, 2020; Porat *et al*., 2022), several questions remain as to the timing and importance of post-transcriptional modifications with respect to other U6 processing steps in organisms like fission yeast and humans.

In addition to 5’ γ-monomethyl phosphate capping enzymes, several other proteins have been linked to U6 processing. These include the La protein, which associates with nascent U6 transcripts through the 3’ uridylate tail (Rinke and Steitz, 1985), and the Lsm2-8 complex, which binds end-matured U6 and remains stably associated through spliceosome assembly (Montemayor *et al*., 2018, 2020; Fu *et al*., 2022). Recent work revealed that mammalian LARP7, a La-Related Protein (LARP) previously linked to MePCE in the context of the 7SK snRNP (He *et al*., 2008), is also involved in post-transcriptional processing of U6. LARP7 promotes 2’-O-methylation of U6 by the methyltransferase fibrillarin, which in turn contributes to splicing fidelity at elevated temperatures in humans and in male germ cells in mice (Hasler *et al*., 2020; Wang *et al*., 2020). Conversely, ciliate and fission yeast LARP7 homologs have been well studied for their roles in telomerase biogenesis (Witkin and Collins, 2004; Collopy *et al*., 2018; Mennie, Moser and Nakamura, 2018; Páez-Moscoso *et al*., 2018; Basu *et al*., 2020; Hu *et al*., 2020). We and others have reported that the *S. pombe* LARP7 protein Pof8 associates with the Bin3/MePCE homolog Bmc1 in the telomerase holoenzyme, that this interaction is important for optimal telomerase activity, and that the link between these proteins is evolutionarily conserved across diverse fungal species (Deragon, 2020; Páez-Moscoso *et al*., 2022; Porat *et al*., 2022). Thus, while much has been learned about MePCE/Bmc1 function in the 7SK snRNP and telomerase, its precise role in U6 biogenesis and function remains unknown.

In this work, we set out to examine the role of Bmc1 in U6 biogenesis and spliceosome function. We have identified a new RNP containing the U6 snRNA and the telomerase components Bmc1, Pof8, and Thc1, and show that this complex is required for wild type levels of 2’-O-methylation in the U6 ISL and U6 snRNP assembly. We also show that while Bmc1’s 5’ capping catalytic activity is not required for its function in promoting 2’-O-methylation of U6, an intact Pof8-Lsm2-8 interaction is. Finally, we show that Bmc1 deletion influences the splicing of some introns under wild-type and heat-shock conditions, consistent with previous work linking human LARP7 to splicing robustness, and that efficient splicing upon heat shock is largely dictated by intronic features including 5’ splice site sequences. Together, these data point towards an intricate network of post-transcriptional processing events that are critical for normal U6 maturation, and provide the first direct evidence for a function of the Bin3/MePCE family in promoting U6 snRNP assembly.

## Results

### Bmc1 forms a U6-containing complex with the telomerase proteins Pof8 and Thc1

Our previous work characterizing Bmc1 as a component of the telomerase holoenzyme also revealed interactions between Bmc1 and various other noncoding RNAs, including the U6 snRNA (Figure S1A) (Porat *et al*., 2022). We therefore tested whether Bmc1 has a role in the biogenesis, stability, or function of these transcripts, and if this function is linked to the Bmc1-interacting telomerase components Pof8 and Thc1. Having already demonstrated that Pof8 is required to recruit Bmc1 to the telomerase RNA TER1 (Porat *et al*., 2022), we determined the protein binding requirements for U6. In contrast to what has been reported for TER1, for which reduced binding to Pof8 persists in the absence of Bmc1 (Páez-Moscoso *et al*., 2022; Porat *et al*., 2022), we found that all three proteins are necessary for an interaction with U6 (Figure 1A, S1B). Our results indicating an interaction between Pof8 and U6 are also consistent with previous work identifying mammalian LARP7 as a U6-interacting protein (Hasler *et al*., 2020; Wang *et al*., 2020), suggesting that LARP7 family members have conserved functions related to U6, in addition to the established evolutionary conservation of LARP7 in telomerase (Collopy *et al*., 2018; Mennie, Moser and Nakamura, 2018; Páez-Moscoso *et al*., 2018).

**Figure 1:**
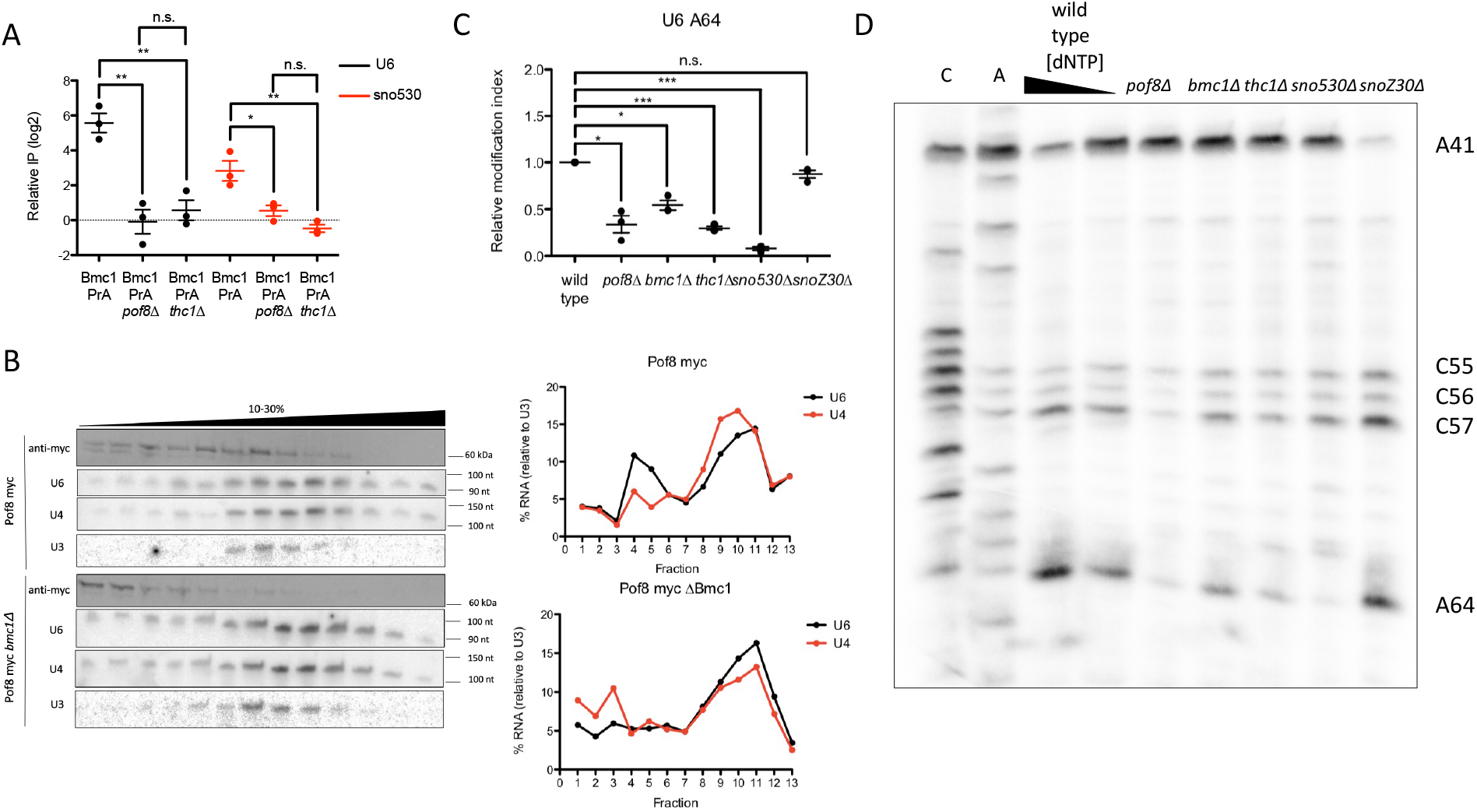
Bmc1, Pof8, and Thc1 guide 2’-O-methylation of U6. A) qRT-PCR of U6 and sno530 in Bmc1 PrA immunoprecipitates, normalized to immunoprecipitation from an untagged strain (mean± standard error, two-tailed unpaired *t* test, \**p*<0.05, \*\**p*<0.01) (n= 3 biological replicates). B) Glycerol gradient sedimentation of myc-tagged Pof8, U4, and U6, and U3 from wild type (Pof8 myc) and *bmc1Δ* strains. U4 and U6 signals were normalized to U3 for calculating relative migration in the gradient. C) Quantification of relative 2’-O-methylation-induced reverse transcriptase stops at A64, compared to a wild type strain (mean± standard error, two-tailed paired *t* test, \**p*<0.05, \*\**p*<0.01, *** *p*<0.001) (n= 3 biological replicates). D) 2’-O-methylation primer extension of U6 at high (1.5 mM) and limiting (0.1 mM) dNTP concentrations. 2’-O-methylated sites are indicated.

As an additional means to confirm U6 snRNP formation, we fractionated native cell extracts on glycerol gradients and compared protein and RNA sedimentation in wild type and knockout yeast strains (Figure 1B, S1C). A substantial fraction of Bmc1 and Pof8 co-migrated with U6, and importantly, co-migration of Pof8 with U6 was impaired upon deletion of Bmc1 (Figure 1B), as well as co-migration of Bmc1 with U6 upon deletion of Pof8 (Figure S1C). We propose that Bmc1, Pof8, and Thc1 associate with U6 simultaneously, with all three proteins required to be present to initiate formation of the Bmc1-containing U6 snRNP. Together, these data point towards the existence of a new U6-containing complex that also shares components with the telomerase holoenzyme, providing a new and surprising link between two seemingly disparate fission yeast noncoding RNA pathways.

### Bmc1, Pof8, and Thc1 promote 2’-O-methylation of U6

To gain further insight into the role of the Bmc1-containing U6 snRNP, we examined our Bmc1 RIP-Seq dataset (Porat *et al*., 2022), which revealed an interaction between Bmc1 and snoZ30, which guides 2’-O-methylation of U6 at position 41 (Zhou *et al*., 2002) (Figure SA, S2A, B). Further supporting the idea that U6 complex formation is contingent on the presence of all three proteins, we observed a loss of snoZ30 binding to Bmc1 upon knockout of any member of the complex (Figure S2A). The observed interaction between Bmc1 and snoZ30, coupled with the well-characterized function of mammalian LARP7 in facilitating snoRNA-guided 2’-O-methylation of U6 by the methyltransferase fibrillarin (Hasler *et al*., 2020; Wang *et al*., 2020) provided initial clues as to the function of this new U6-containing snRNP. To determine if Bmc1, Pof8, and Thc1 influence 2’-O-methylation, we mapped U6 2’-O-methylation sites by performing primer extensions at low dNTP concentrations (Zhou *et al*., 2002). Although snoZ30 is the sole annotated U6-modifying snoRNA in fission yeast (Zhou *et al*., 2002), several other 2’-O-methylated sites have been identified in U6, including A64 (Gu *et al*., 1996). Deletion of Bmc1, Pof8, and Thc1 resulted in no observable changes in 2’-O-methylation at the snoZ30-modified A41, but we did detect a reproducible decrease in modification at several other sites, most notably A64 (Figure 1C, D, Figure S3).

Initial attempts at identification of the U6 A64-methylating snoRNA using box C/D snoRNA consensus sequences and base pairing rules (Lowe and Eddy, 1999) yielded no other obvious snoRNA candidates, so we instead turned to our Bmc1 RIP-Seq dataset in the hope we might identify novel snoRNAs (Figure S1A). The uncharacterized fission yeast noncoding RNA, SPNCRNA.530 (henceforth referred to as sno530), contains a D box, a putative C box one nucleotide different from the C box consensus motif, and a region with 12 nucleotides of complementarity with U6, with a single non-Watson Crick base pair (Figure S2C). It is also noteworthy that the predicted secondary structure of sno530 does not position the C and D boxes flanking a hairpin, as is common for canonical box C/D snoRNAs (Figure S2C). We validated the interaction between Bmc1 and sno530 by RNP immunoprecipitation/qPCR and showed that much like snoZ30 and U6, this interaction is dependent on the presence of the assembled Bmc1-Pof8-Thc1 complex (Figure 1A). Deletion of snoZ30 and sno530 resulted in a loss of 2’-O-methylation at A41 and A64, respectively, suggesting that sno530 is indeed the A64 U6-modifying snoRNA (Figure 1C, D, Figure S3). We obtained similar results using a complementary method that exploits the tendency for 2’-O-methylations to block RNase H cleavage following the annealing of a chimeric DNA-2’-O-methylated RNA oligo targeting the suspected 2’-O-methylated site (Yu, Shu and Steitz, 1997; Calo *et al*., 2015) (Figure S4A). This also served to provide evidence for 2’-O-methylation at C57, suggesting that it, too, is another site in U6 whose modification is similarly guided by Bmc1 and Pof8 (Figure S4B). While our knockout studies unambiguously identify sno530 as the A64 U6-modifying snoRNA, the unusual sequence and architecture of sno530 relative to snoZ30 is more reminiscent of the divergent box C’/D’ motifs that stimulate rRNA 2’-O-methylation by providing additional regions of complementarity surrounding the methylated site (Van Nues *et al*., 2011; van Nues and Watkins, 2017).

### Bmc1, Pof8, and Thc1 are involved in U6 snRNP assembly

We then wondered how disruption of the Bmc1-containing U6 snRNP might impact spliceosome assembly. We observed small differences in U6 sedimentation in glycerol gradients, with a less abundant, lighter sedimenting U6-containing fraction appearing in addition to co-sedimentation with U4 (compare U6 and U4 in lanes 3-6 in Figure 1B and S1C), suggesting a U6-containing complex without U4. Consistent with this, we note that the migration of Pof8, Bmc1, and Thc1 does not fully overlap with U4/U6 in the gradient, but is rather shifted towards lighter fractions, arguing against the complete inclusion of the Bmc1-Pof8-Thc1 complex in the U4/U6 di-snRNP (Figure 1B, S1C). As the lighter sedimenting U6 species is not evident in Bmc1 and Pof8 KO strains (compare U6 and U4 in lanes 3-6 in Figure 1B and S1C relative to Bmc1 and Pof8 KO strains), we hypothesized that Bmc1, Pof8, and Thc1 interact with U6 before the U4/U6 di-snRNP.

To obtain clearer resolution of distinct U6-containing complexes, we ran cell extracts on native gels and analyzed spliceosomal RNAs by northern blotting. We observed a single, prominent band for all spliceosomal RNAs except U6, which migrated as 2 distinct complexes (Figure 2A). We could assign the higher molecular weight complex, which comigrates with U4 but not U2 or U5, as the U4/U6 di-snRNP. The U4/U6 di-snRNP, as well as other spliceosomal snRNPs and the non-spliceosomal U3 snRNP, showed no change in relative intensity or migration upon deletion of Bmc1, Pof8, Thc1, or sno530. However, we did observe a significant and reproducible decrease in the intensity of the lower molecular weight U6-containing snRNP upon deletion of Bmc1, Pof8, and Thc1 (Figure 2A, B), consistent with this band representing the Bmc1-containing U6 snRNP. The persistence of this complex upon loss of sno530 suggests that complex formation is not reliant on the ability to modify U6 at A64. Although U6 and sno530 are associated with Bmc1 (Figure 1A), sno530 is therefore not required for the stability of the U6 snRNP observed in native gels.

**Figure 2:**
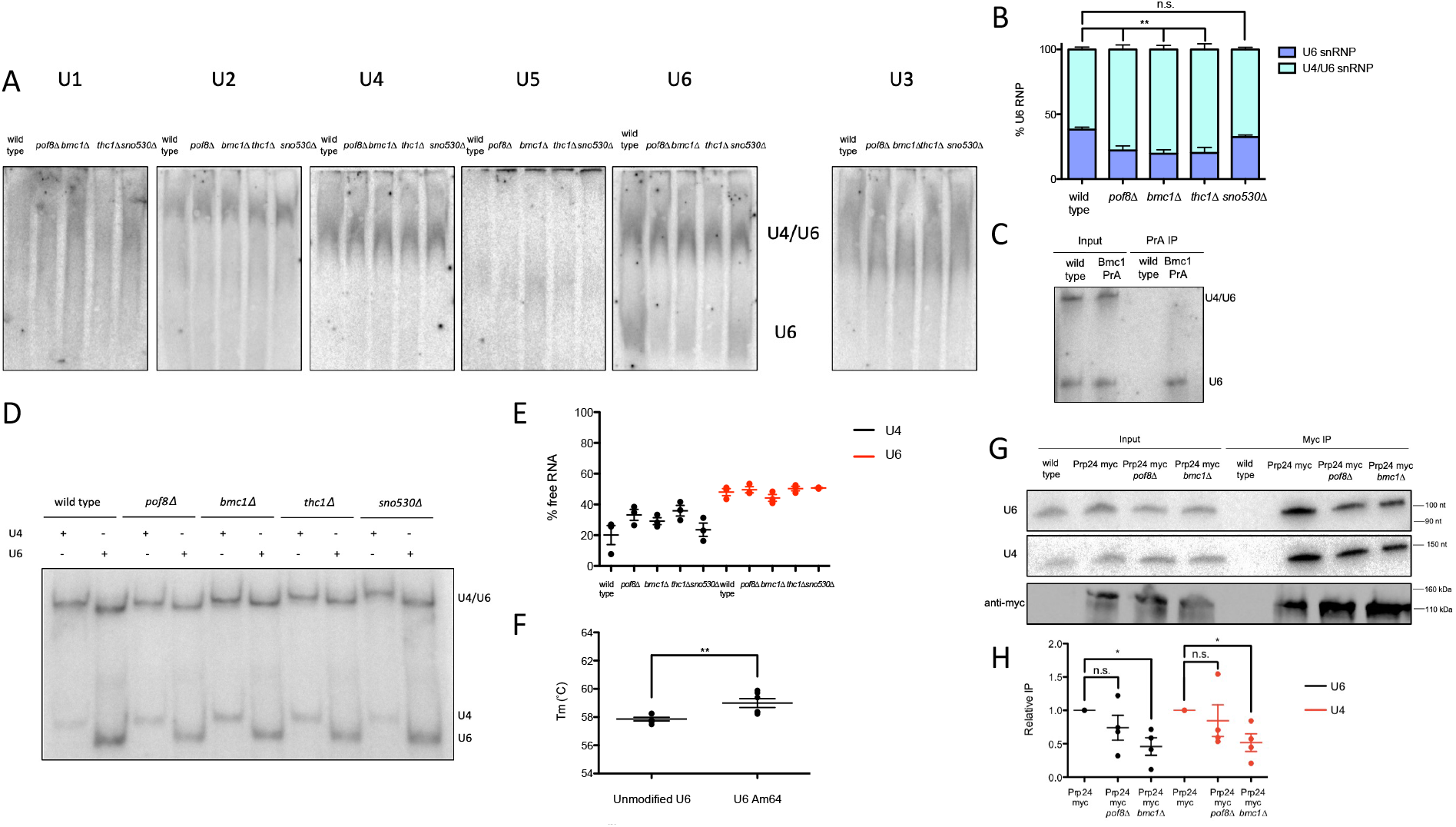
Bmc1, Pof8, and Thc1 promote U6 snRNP assembly. A) Native northern blot analysis of spliceosomal and non-spliceosomal (U3) snRNPs from native yeast cell extracts. B) Quantification of U6-containing snRNPs from wild type and knockout yeast cell extracts (mean± standard error, two-tailed unpaired *t* test, \*\**p*<0.01) (n= 4 biological replicates). C) Native northern blot analysis of total and Bmc1-immunoprecipitated U6. D) Solution hybridization of U4/U6 pairing in wild type and knockout yeast strains using radiolabeled probes targeting the 5’ end of U4 and 3’ end of U6. E) Quantification of U4/U6 pairing from solution hybridization assay, expressed as the fraction of non-duplexed U4 and U6 (“free RNA”) (mean± standard error) (n= 3 biological replicates). F) Tm values from UV melt curve analysis of U4/U6 pairing with unmodified and A64-2’-O-methylated U6 oligos (mean± standard error, two-tailed unpaired *t* test, \*\**p*<0.01) (n= 6 technical replicates). G) Northern and western blot analysis of U4, U6, and myc-tagged Prp24 from total cell extracts and myc-immunoprecipitates. H) Quantification of Prp24-immunoprecipitated U4 and U6, relative to Prp24 myc (mean± standard error, two-tailed paired *t* test, \**p*<0.05) (n= 4 biological replicates).

To understand when Bmc1 interacts with U6 with respect to spliceosome formation, we extracted Bmc1 immunoprecipitated RNPs under native conditions and ran total and Bmc1-associated RNA on native gels. Bmc1-bound U6 did not co-migrate with the U4/U6 di-snRNP (Figure 2C), suggesting that Bmc1 interacts with U6 outside of the U4/U6 di-snRNP, in line with the Bmc1/Pof8/Thc1-associated mono U6 snRNP band (2A, B). Based on these results, we hypothesize that the Bmc1-containing U6 complex, which promotes 5’ capping and 2’-O-methylation, forms upstream of the U4/U6 di-snRNP.

Native fission yeast cell extracts do not form detectable amounts of the U4/U6.U5 tri-snRNP (Huang, Vilardell and Query, 2002; Chen *et al*., 2014), so we focused our further efforts on examining U4/U6 base pairing by performing a solution hybridization assay on cold phenol-extracted total RNA to maintain U4/U6 base pairing (Figure 2D). This differs from native spliceosomal snRNP gels (Figure 2A) in that it only assesses RNA-RNA interactions, without changes in mobility due to protein binding. We detected minor defects in U4/U6 assembly upon Bmc1, Pof8, or Thc1 deletion, as measured by the increase in “free” U4 relative to U4 complexed in the di-snRNP, although the increase in the fraction of free U4 did not reach statistical significance (Figure 2D, E). Consistent with the increase in free U4 in the knockout strains, glycerol gradients revealed an increase in lighter sedimenting U4 in the knockout strains (Figure 1B, S1B, compare lanes 1-3 in wild type versus knockouts). Although U6 is in excess over U4, the increase of free U4 in the knockouts suggests that the absence of Bmc1, Pof8, and Thc1 may result in a non-functional, alternate pathway for U4 that does not involve U4/U6 di-snRNP formation. The lack of U4/U6 pairing defects upon the loss of sno530 further suggests that it is largely the Bmc1-Pof8-Thc1 protein complex dictating U4/U6 pairing, not the single A64 2’-O-methylation. Still, UV melt analysis of the U6-interacting region of U4 and the U6 internal stem loop (ISL), with or without 2’-O-methylation of A64, revealed a slight increase in U4-U6 duplex stability with 2’-O-methylation, consistent with previous findings reporting on the stabilizing properties of 2’-O-methylation on RNA duplex formation (Prusiner, Yathindra and Sundaralingam, 1974; Kawai *et al*., 1992; Abou Assi *et al*., 2021) (Figure 2F).

Subsequent steps in spliceosome formation involve unwinding of the U6 ISL and forming base pairing between U4 and U6, both of which are promoted by the U4/U6 di-snRNP assembly factor Prp24 (Ghetti, Company and Abelson, 1995; Martin-Tumasz *et al*., 2011). We generated an endogenously tagged Prp24 strain and assessed the interaction between Prp24 and U4 and U6. Pof8 and Bmc1 deletion resulted in a decreased interaction between Prp24 and U4 and U6, suggesting that Bmc1 and Pof8 promote the association of U4 and U6 with Prp24, which in turn may promote the formation of the U4/U6 di-snRNP (Figure 2G, H). In sum, our results are consistent with the existence of a Bmc1/Pof8/Thc1-containing U6 snRNP, with Bmc1/Pof8/Thc1 dissociating from U6 during establishment of the U4/U6 di-snRNP.

### Bmc1 5’ capping catalytic activity is not required for promoting 2’-O-methylation of U6

With previous studies indicating that Bmc1 5’ γ-phosphate methyltransferase catalytic activity is dispensable for telomerase activity (Páez-Moscoso *et al*., 2022), we assayed a combination of previously described and newly constructed putative Bmc1 catalytic mutants for the ability to promote U6 2’-O-methylation. We mutated residues that are both highly conserved between Bmc1 and human MePCE, and well-positioned in structure predictions to interact with the methyltransferase byproduct SAH (Figure 3A). HA-tagged Bmc1 mutants were transformed into a Bmc1 knockout yeast strain and profiled for U6 2’-O-methylation as above (Figure 3B, C). While the Bmc1 mutants were more lowly expressed than wild type Bmc1, some mutants still promoted 2’-O-methylation to a greater extent than the empty vector (Figure 3C). Further, normalization of relative 2’-O-methylation levels to Bmc1 expression confirmed a statistically significant increase in 2’-O-methylation for all Bmc1 mutants compared to the empty vector (Figure 3D). This suggests that, as in telomerase, Bmc1 5’ γ-phosphate methyltransferase catalytic activity is not critical for its function in U6 2’-O-methylation.

**Figure 3:**
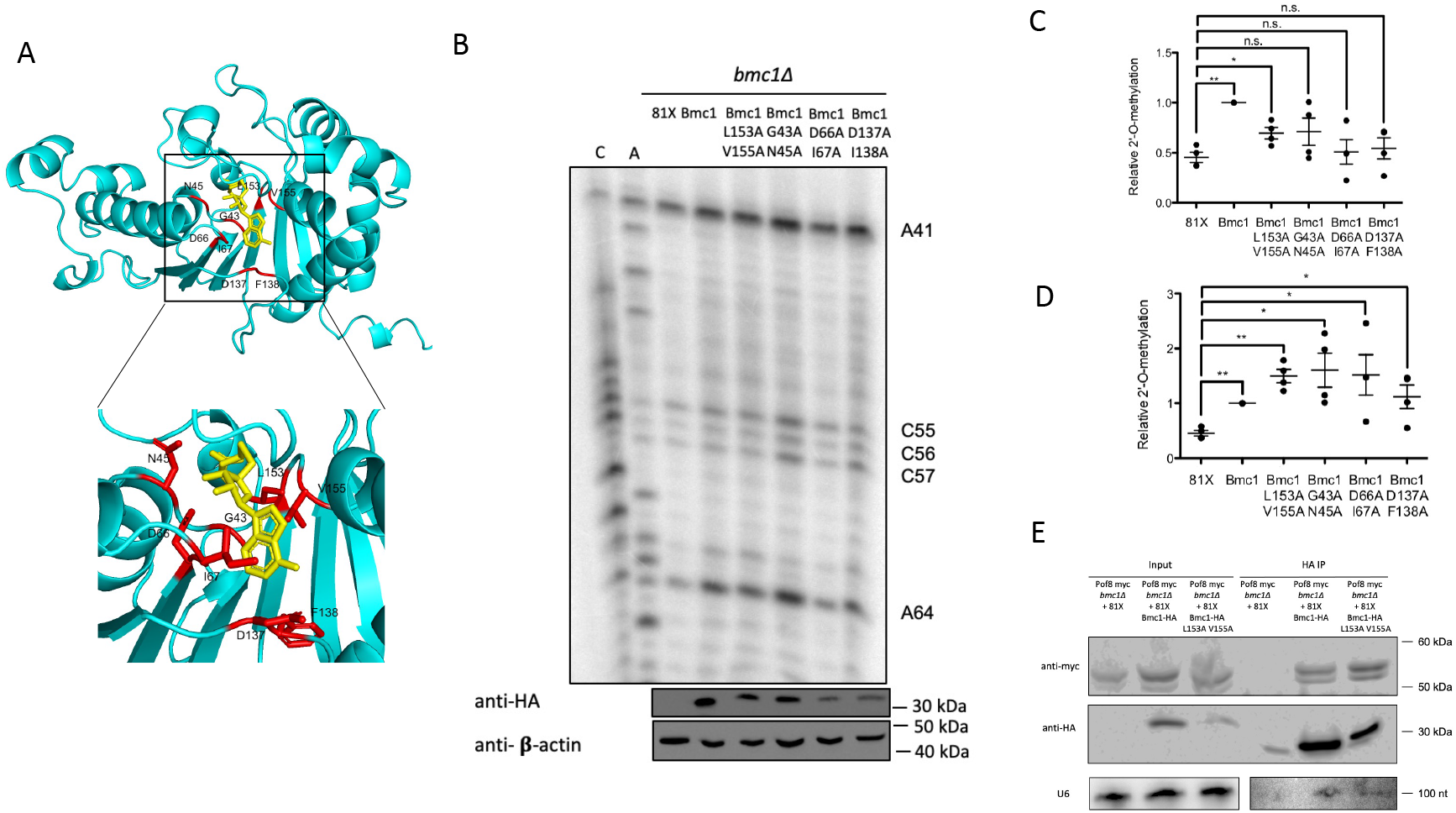
Bmc1 catalytic activity is not a requirement for 2’-O-methylation of U6. A) AlphaFold (Jumper *et al*., 2021) structure prediction of Bmc1 aligned to the SAH-bound (yellow) catalytic domain of MePCE (PDB 6DCB) (Yang *et al*., 2019) with mutations indicated in red. Inset: side chain interactions with SAH. B) U6 2’-O-methylation primer extension in *bmc1Δ* cells transformed with the indicated plasmid. 2’-O-methylated sites are indicated. Western blots for Bmc1-HA expression and b-actin are indicated below. C) Quantification of relative 2’-O-methylation-induced reverse transcriptase stops at A64, compared to wild type Bmc1-HA (mean± standard error, two-tailed paired *t* test, \**p*<0.05, \*\**p*<0.01) (n= 4 biological replicates). D) Quantification of relative 2’-O-methylation-induced reverse transcriptase stops at A64, compared to wild type Bmc1-HA, normalized to average Bmc1-HA expression relative to b-actin (mean± standard error, two-tailed paired *t* test, \**p*<0.05, \*\**p*<0.01) (n= 4 biological replicates). E) Western blot and northern blot analysis of co-immunoprecipitation of HA-tagged Bmc1, myc-tagged Pof8 and U6.

For further characterization of Bmc1 catalytic mutants, we chose the Bmc1 L153A V155A mutant, which showed the highest expression across biological replicates. As measured by co-immunoprecipitation, L153A V155A still interacted with Pof8, suggesting that catalytic activity is also not required for complex formation (Figure 3E). Further, L153A V155A interacted with U6, indicating that 5’ γ-phosphate methyltransferase catalytic activity is not required for U6 binding (Figure 3E).

### The xRRM and Pof8-Lsm2-8 interaction are important determinants for U6 2’-O-methylation

As an established member of the LARP7 family of proteins, the protein-interacting and RNA binding domains of Pof8 have been well-characterized in the context of the telomerase RNP (Collopy *et al*., 2018; Mennie, Moser and Nakamura, 2018; Páez-Moscoso *et al*., 2018; Basu *et al*., 2020; Hu *et al*., 2020). Pof8 contains a divergent La motif that lacks the conserved uridylate-binding residues typically seen in LARP7 proteins (Páez-Moscoso *et al*., 2018), so its interaction with the telomerase RNA TER1 is mediated by the RRM1, xRRM, and the N-terminal region that makes direct protein-protein contacts to Lsm2-8, which in turn binds the uridylate-rich 3’ end of TER1 (Collopy *et al*., 2018; Mennie, Moser and Nakamura, 2018; Páez-Moscoso *et al*., 2018; Hu *et al*., 2020). As mutations to these regions have been shown to impair Pof8 binding to TER1 and telomere length homeostasis, we looked at the impact of these same mutations on U6 2’-O-methylation (Figure 4A). In contrast to what has been observed for TER1, where both RRMs are important for binding, only mutations to the xRRM and the Lsm2-8 binding region caused a significant reduction in 2’-O-methylation at A64 (Figure 4B, C).

**Figure 4:**
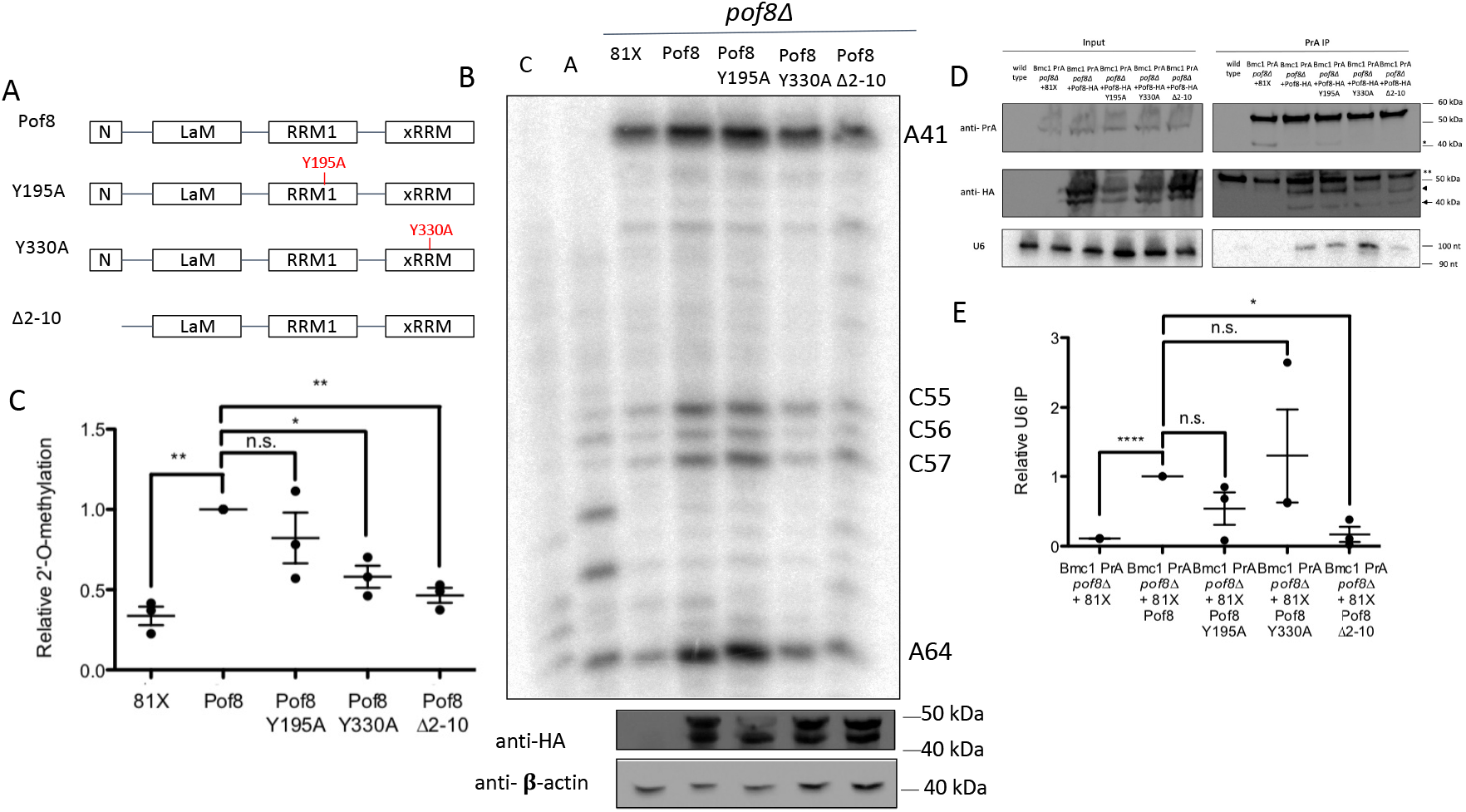
The xRRM and Lsm2-8-binding surface are important in Pof8-mediated 2’-O-methylation of U6. A) Schematic of Pof8 domains and mutants used in this study. N= N-terminal domain, LaM= La motif, RRM1= RNA Recognition Motif 1, xRRM= extended RNA Recognition Motif. B) U6 2’-O-methylation primer extension in *pof8Δ* cells transformed with the indicated plasmid. 2’-O-methylated sites are indicated. Western blots for Pof8-HA expression and b-actin are indicated below. C) Quantification of relative 2’-O-methylation-induced reverse transcriptase stops at A64, compared to wild type Pof8-HA (mean± standard error, two-tailed paired *t* test, \**p*<0.05, \*\**p*<0.01) (n= 3 biological replicates). D) Northern and western blot analysis of U6, PrA-tagged Bmc1, and HA-tagged Pof8 from total cell extracts and PrA-immunoprecipitates. *Bmc1 PrA cleavage products, **An additional band cross-reacting with the antibody, arrows indicate Pof8 HA. E) Quantification of Bmc1-immunoprecipitated U6, relative to the Pof8-HA-expressing strain (mean± standard error, two-tailed paired *t* test, \**p*<0.05, \*\**p*<0.01, ****p<0.0001) (n= 3 biological replicates).

To further understand the molecular basis for the drop in 2’-O-methylation, we immunoprecipitated Bmc1 in a Pof8 knockout strain re-expressing the Pof8 mutants (Figure 4D, E). Bmc1 co-immunoprecipitated all Pof8 mutants, suggesting that the 2’-O-methylation defect is not due to complete disruption of the Bmc1-Pof8 interaction (Figure 4D). The Bmc1-U6 interaction, which is dependent on the presence of Pof8 (Figure 1A), was almost completely lost in the Lsm2-8-binding mutant (Δ2-10), suggesting that its 2’-O-methylation defect may be due to a loss in U6 association with the Bmc1-Pof8-Thc1 complex (Figure 4D, E). Surprisingly, we detected no loss in U6 binding with the xRRM mutant, indicating that while U6 still interacts with the Bmc1-Pof8-Thc1 snRNP in the context of the xRRM mutant, the xRRM may have another function in facilitating U6 2’-O-methylation (Figure 4D, E), potentially through the binding of the snoRNA, as has been suggested for human LARP7 (Hasler *et al*., 2020). Alternatively, as the xRRM in the ciliate LARP7 protein p65 has been suggested to possess RNA chaperone activity to remodel the ciliate telomerase RNA (Singh *et al*., 2012; Singh, Choi and Feigon, 2013), it is tempting to speculate that similar xRRM-mediated RNA chaperone activity may play a role in correctly positioning U6 for 2’-O-methylation.

### Bmc1 deletion has a minor effect on intron retention

Having observed Bmc1-dependent defects in U6 2’-O-methylation and U6 snRNP assembly, we tested the effects of Bmc1 deletion on pre-mRNA splicing. To that end, we performed shortread, paired-end sequencing on RNA extracted from wild type and Bmc1 knockout yeast strains and quantified intron retention as a proxy for splicing (Lorenzi *et al*., 2021; Schärfen *et al*., 2022). We also measured intron retention in wild type and knockout cells heat shocked for 15 minutes at 42°C, which has been shown to impact splicing in fission yeast (Awan, Manfredo and Pleiss, 2013). We observed significant increases in intron retention following heat shock, similar to what has been reported in mammalian cells (Shalgi *et al*., 2014) (Figure S5A, B). Although we observed slight increases in intron retention in Bmc1 knockout cells compared to wild type cells, very few of these splicing events at 32°C passed our significance cut-off, and no splicing events at 42°C were statistically significant (Figure S5C, D), although this may be due to greater sample to sample variability across our triplicate replicates for this data set (Figure S6). Still, as mean intron retention values indeed showed an increase upon Bmc1 deletion (Figure 5A), we chose several representative intron retention events to validate with semi-quantitative RT-PCR (one of which, intron 1 of pud1, displayed a statistically significant increase upon Bmc1 deletion at 32°C in our RNA Seq dataset). We observed an increase in intron retention following heat shock, and confirmed their further impaired splicing in the context of the Bmc1 deletion (Figure 5B, S7). Conversely, a ribosomal protein gene, which have been reported to be efficiently spliced relative to non-ribosomal protein genes in budding yeast (Barrass *et al*., 2015; Gildea, Dwyer and Pleiss, 2022), and did not show heat shock or Bmc1 associated changes in our RNA-Seq data set, was confirmed to have no changes in intron retention in response to heat shock or Bmc1 deletion (Figure 5B, S7). We note that these validated Bmc-1 affected introns have higher than average intron retention rates in normal cells and as such, may not be representative of the average splicing event. Together, these data indicate that Bmc1 does not have a major effect on pre-mRNA splicing, but that Bmc1 likely contributes to splicing robustness, similar to what has been described for mammalian LARP7 deletion in human cells (Hasler *et al*., 2020; Wang *et al*., 2020).

**Figure 5:**
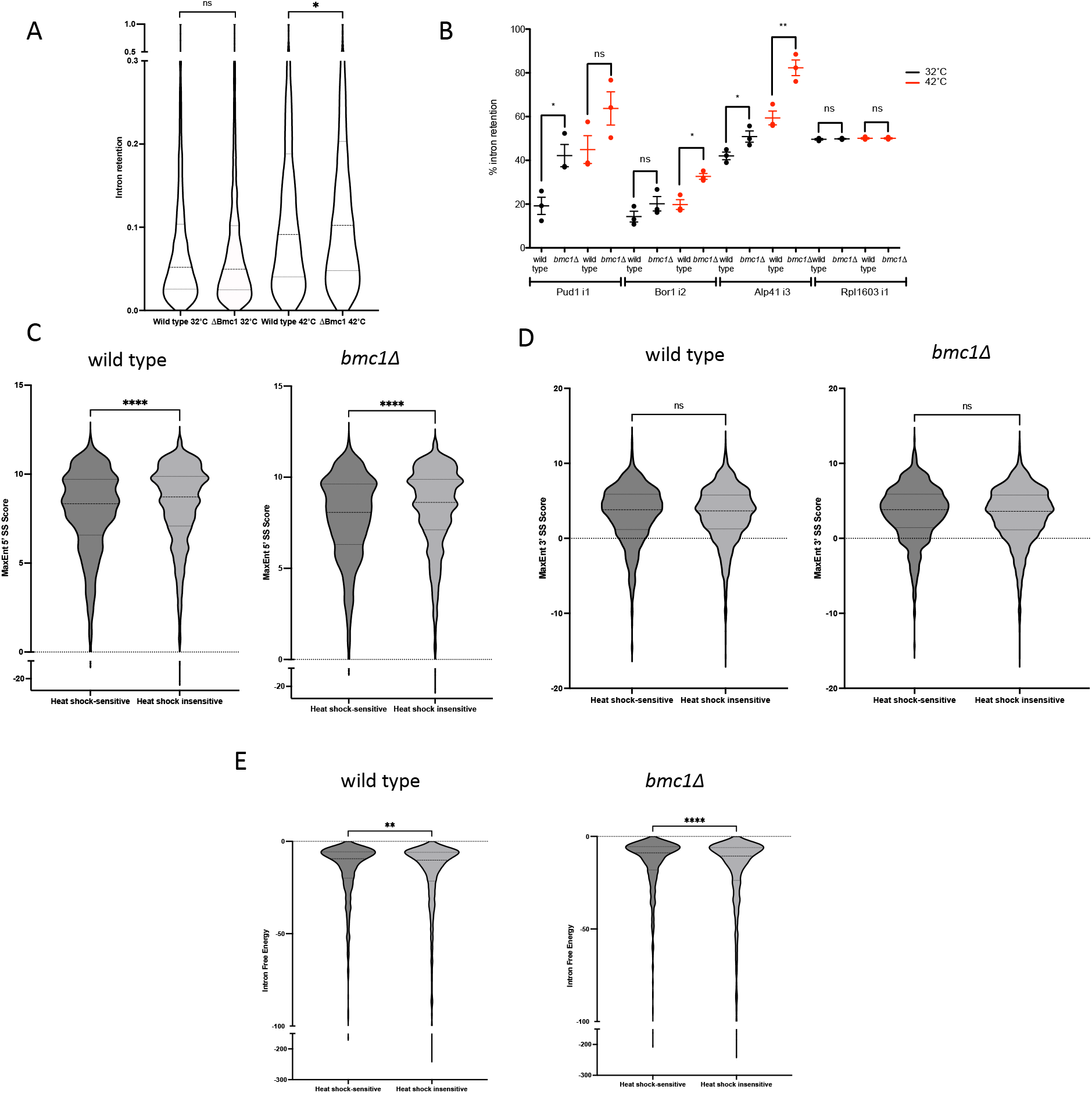
Bmc1 deletion leads to minor splicing defects. A) Average intron retention from 3 biological replicates for wild type and *bmc1Δ* strains grown at 32°C or heat shocked at 42°C (two-tailed unpaired t-test with Welch’s correlation, * *p*<0.05). B) Semi-quantitative RT-PCR in wild type and *bmc1Δ* strains grown at 32°C or heat shocked at 42°C for intron 1 of *pud1*, intron 2 of *bor1*, intron 3 of *alp41*, and intron 1 of *rpl1603* (mean± standard error, two-tailed unpaired *t* test, \**p*<0.05, \*\**p*<0.01) (n= 3 biological replicates). C-E) Comparison of 5’ splice site scores (C), 3’ splice site scores (D), and intron minimum free energy (kcal/mol) (E) for heat shock-sensitive and insensitive introns in wild type and *bmc1Δ* cells. Heat shock-sensitive introns were classified as introns that exhibited a greater than 2-fold increase in intron retention following heat shock and a false discovery rate less than 0.05. Remaining introns are classified as heat shock insensitive. Only introns with greater than 4 reads supporting splicing in all biological replicates were included (two-tailed unpaired t-test with Welch’s correlation, \*\**p*<0.01, \*\*\*\**p*<0.0001).

We also examined other factors that could contribute to heat shock-sensitive splicing defects. We compared intronic features between heat shock-sensitive introns, classified as introns exhibiting a greater than 2-fold increase in intron retention upon heat shock and a false discovery rate less than 0.05, and remaining introns (heat shock insensitive) (Figure 5C-E, S5A, B). In line with previous data indicating that intron retention in fission yeast can arise from a weak 5’ splice site (Melangath *et al*., 2017), we noted that heat shock-sensitive introns have a weaker 5’ splice site score in the context of a wild type and *bmc1Δ* strain, with no significant differences in 3’ splice site scores (Figure 5C,D). Additionally, heat shock-sensitive introns displayed lower minimum free energy, indicative of a link between intron structure and splicing changes in response to heat shock (Figure 5E).

## Discussion

### Conserved functions for LARP7 family proteins in splicing and U6 2’-O-methylation

This work represents the first report of an MePCE homolog with a role in splicing and spliceosome assembly, beyond 5’ methyl phosphate cap addition of U6. In our efforts to investigate functions for Bmc1 beyond telomerase, we revealed an unanticipated overlap between components of the yeast telomerase holoenzyme and a U6-containing snRNP. While it is surprising that Bmc1, Pof8, Thc1, and Lsm2-8 interact with 2 very distinct non-coding RNAs produced by different polymerases, both RNAs possess uridylate-rich sequences recognized by Lsm2-8 and highly structured regions, including stem loops in U6 and pseudoknots in telomerase, that act as scaffolds to recruit other RNP components. These common features may provide an explanation as to why these divergent RNAs share a common set of protein binding partners. Such RNP plasticity is not unique to fission yeast telomerase and U6, but may represent a shared feature of LARP7 and MePCE family proteins. Mammalian LARP7 and MePCE are particularly well-studied for their roles in capping and stabilizing the 7SK snRNP (Jeronimo *et al*., 2007; He *et al*., 2008; Krueger *et al*., 2008; Markert *et al*., 2008), transcriptional control through DDX21 (Calo *et al*., 2015), directing U6 modification (Hasler *et al*., 2020; Wang *et al*., 2020), and snRNP assembly through the SMN complex (Ji *et al*., 2021). Thus, continuing to study the RNA interactome of MePCE and LARP7 homologs across species will likely yield additional insight into how these proteins associate with and influence various classes of non-coding RNAs. It is also possible that Bmc1, Pof8, and Thc1 interactions with U6 are mediated entirely by direct interactions with Lsm2-8, which in turn directly contacts U6, much like Prp24 interacts with U6 by directly binding Lsm2-8 (Rader and Guthrie, 2002). Future structural and biochemical studies will lend insight into the protein-protein and protein-RNA interactions that cooperate to form the U6 snRNP.

Several fungal species, including *S. cerevisiae*, lack both a LARP7 and MePCE homolog (Deragon, 2020; Porat *et al*., 2022). As deletion or depletion of LARP7/Pof8 or MePCE/Bmc1 does not influence U6 stability in species where this has been investigated (Jeronimo *et al*., 2007; Hasler *et al*., 2020; Wang *et al*., 2020; Páez-Moscoso *et al*., 2022; Porat *et al*., 2022), the function of LARP7 and MePCE family members in U6 biogenesis and function has remained unclear. This work expands our understanding of the evolutionary conservation of LARP7 family members, with shared or unique functions relating to the telomerase, U6, and 7SK RNAs, depending on the species under investigation (Figure 6A). Further links can be drawn between the functional consequences of LARP7- and Pof8-mediated promotion of U6 2’-O-methylation. LARP7 or Pof8 deletion and the subsequent decrease in 2’-O-methylation of U6 results in no functional consequences under standard physiological conditions, but becomes important for maintaining splicing fidelity under heat stress, in the case of human LARP7, and male germ cells in mice (Hasler *et al*., 2020; Wang *et al*., 2020). As the loss of A64 modification alone results in no changes to U6 snRNP assembly, compared to the changes observed upon Bmc1, Pof8, and Thc1 deletion, we anticipate that it is either a combination of the loss of several 2’-O-methylations or the loss of the Bmc1-Pof8-Thc1 U6 snRNP that leads to the slight increase in intron retention at elevated temperatures upon Bmc1 deletion (Figure 3). Future studies aimed at teasing apart this mechanism in mammalian and yeast cells will provide additional insight into the intertwining role of RNA modifications and RNP biogenesis complexes in spliceosome assembly.

**Figure 6:**
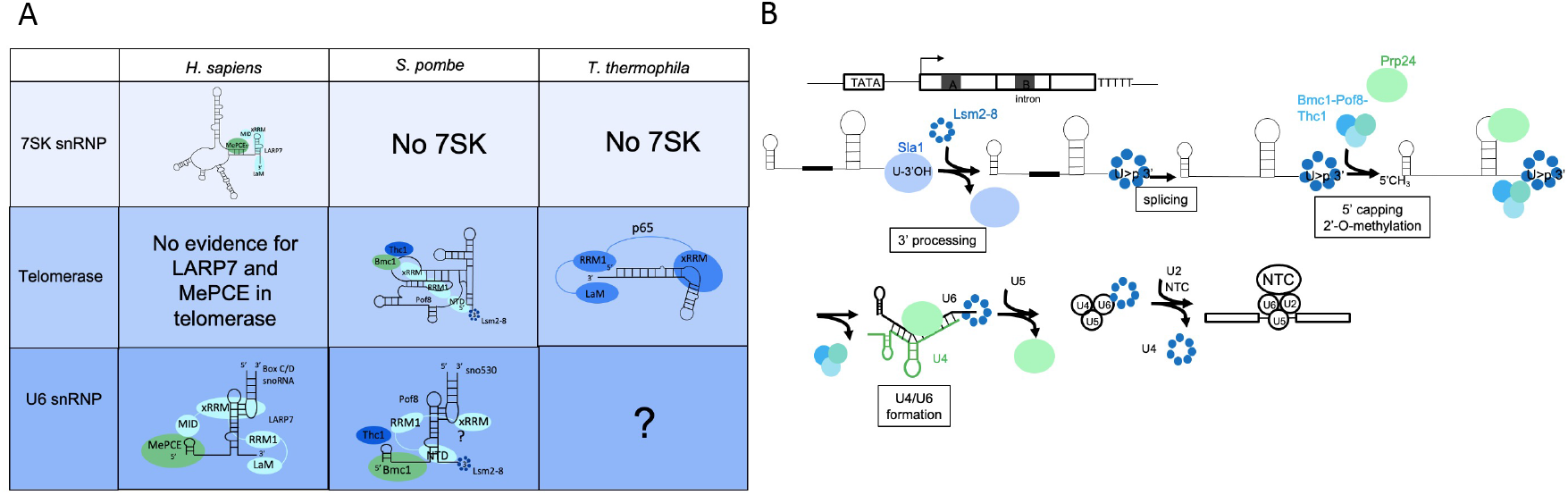
Evolutionary convergence and divergence of Bmc1/MePCE and Pof8/LARP7 in noncoding RNA processing. A) Summary of Bmc1/MePCE and Pof8/LARP7/p65 functions in the 7SK snRNP, telomerase holoenzyme, and U6 snRNP. LaM= La motif, RRM1= RNA Recognition Motif 1, xRRM= extended RNA Recognition Motif, NTD= N-terminal domain (Lsm2-8-interacting region), MID= MePCE-Interacting Domain. The existence and composition of a U6 snRNP in *T. thermophila* is currently unknown. B) Schematic of the U6 biogenesis pathway in fission yeast. NTC= NineTeen Complex.

Another question that emerges from this study concerns the minor splicing defects observed in Bmc1 knockout cells at elevated temperatures. Previous studies on mammalian LARP7 proposed that LARP7-guided 2’-O-methylation of U6 is not an important factor for splicing as a whole, but rather contributes to splicing robustness (Hasler *et al*., 2020; Wang *et al*., 2020). Although recent reports indicate that alternative splicing in fission yeast may be more widespread than previously thought (Awan, Manfredo and Pleiss, 2013; Kuang, Boeke and Canzar, 2017; Montañés *et al*., 2022), splicing complexity in fission yeast is still less than that observed in mammalian cells, which may explain why we do not observe any drastic splicing changes upon Bmc1 deletion. It remains to be determined whether Bmc1 affects other aspects of splicing that have not been tested here, such as splicing efficiency and fidelity, or whether Bmc1-associated defects in splicing might be greater under different stresses. In addition, Bmc1 promotes 2’-O-methylation in the internal stem loop of U6, which does not base pair with the 5’ or 3’ splice site. Thus, modulation of ISL modifications might not be expected to manifest as a robust splicing defect. This is in contrast to what has been reported for the loss of m^6^A in *S. pombe* U6, where affected introns are enriched for an adenosine at the fourth position of the intron, which directly base pairs with the m^6^A (Ishigami *et al*., 2021).

### Emerging importance of the xRRM in RNA folding and function

Fission yeast, possessing a LARP7 homolog that functions in telomerase like its ciliate counterpart (Collopy *et al*., 2018; Mennie, Moser and Nakamura, 2018; Páez-Moscoso *et al*., 2018, 2022; Basu *et al*., 2020; Hu *et al*., 2020; Porat *et al*., 2022), and U6 2’-O-methylation in an analogous manner to its mammalian homologs, may represent an evolutionary intermediate bridging RNA binding proteins between ciliates and mammals. The 7SK snRNA, which has only been found in animals (Marz *et al*., 2009) likely arose independently from the more widely distributed LARP7 and MePCE, suggesting the need for continued studies into 7SK-independent functions for LARP7 and MePCE. Of note, the conservation of the xRRM between fungal, mammalian, and ciliate LARP7 proteins, rather than the La motif (Collopy *et al*., 2018; Mennie, Moser and Nakamura, 2018; Páez-Moscoso *et al*., 2018; Deragon, 2020; Porat *et al*., 2022) may provide a reason explaining the diverse RNA substrates bound by LARP7 homologs, compared to the more well-conserved classes of RNA binding partners of other LARPs across species (Deragon, 2020). xRRM-mediated binding to structured stem loops like the telomerase RNA pseudoknot (Hu *et al*., 2020), SL4 of 7SK (Eichhorn *et al*., 2018), and U6-modifying snoRNAs (Hasler *et al*., 2020) may be a better determinant than 3’ terminal uridylate stretches for predicting LARP7 binding. The importance of the xRRM in the biogenesis and stability of telomerase RNA, 7SK, and U6 may be linked to its RNA chaperone activity, which has been proposed to have a role in promoting RNA folding (Singh *et al*., 2012; Singh, Choi and Feigon, 2013). Our finding that mutation of the xRRM of Pof8 impairs 2’-O-methylation of U6 without disrupting U6 binding (Figure 5D) may provide further evidence that the xRRM has functions beyond U6 binding and raises additional questions as to the mechanism by which RNA chaperones can coordinate snoRNA and target RNA binding to carry out efficient 2’-O-methylation. Importantly, xRRM chaperone activity is not limited to LARP7 family proteins, as the RRM2/xRRM of the human La protein has also been shown to promote RNA folding (Kucera, Hodsdon and Wolin, 2011; Naeeni, Conte and Bayfield, 2012; Brown *et al*., 2016).

### New insights into U6 biogenesis in fission yeast

This work also sheds light on the timing of U6 biogenesis steps in fission yeast (Figure 6B). We have previously shown that Lsm2-8 interacts with both mature and intron-containing U6, suggesting that intron removal occurs after 3’ end processing and the switch from La to Lsm2-8 (Montemayor *et al*., 2020; Porat *et al*., 2022). Conversely, Bmc1 and Pof8 interact solely with the spliced form of U6 (Porat *et al*., 2022). This, coupled with our finding that the Lsm2-8-interacting region of Pof8 is required for the Bmc1-U6 interaction (Figure 5D, E), indicates that Lsm2-8 binding occurs prior to splicing and recruitment of the Bmc1-Pof8-Thc1 complex. Our data indicating that Bmc1 co-purifies with U6-modifying snoRNAs (Figure 1 and Figure S2) suggests that U6 then undergoes 5’ capping by Bmc1 and 2’-O-methylation, prior to Bmc1-Pof8-Thc1 dissociation from U6 and U4/U6 di-snRNP assembly mediated by Prp24. Since deletion of Bmc1 or Pof8 results in decreased association of Prp24 with U4 and U6 (Figure 2), the Bmc1-Pof8-Thc1 complex may play a role in the handoff to Prp24. This role may be mediated by xRRM-linked chaperone activity that remodels U6 to better position it to interact with Prp24 and U4. Our finding of a new U6 biogenesis complex thus adds another layer of regulation to spliceosome assembly. Still, it remains unknown whether Bmc1, Pof8, and Thc1 only interact with U6 during its biogenesis, or re-associate with U6 when it is reassembled into the U4/U6 di-snRNP for subsequent rounds of splicing catalysis. Notably, our finding of a mono-U6 snRNP containing Bmc1 and Pof8 that promotes internal modifications of U6 is consistent with earlier reports of the human m^6^A methyltransferase METTL16 present in a mono-U6 snRNP with MePCE and LARP7 (Warda *et al*., 2017). Since mammalian U6 also undergoes 5’ methyl phosphate capping by MePCE and LARP7-mediated 2’-O-methylation, it will be interesting to examine the interplay between MePCE, LARP7, and METTL16, and how these factors may function in promoting the formation of the U4/U6 di-snRNP in higher systems.

Taken together, this work adds to the growing body of literature on the catalytic-independent functions of RNA modification enzymes (reviewed in (Porat, Kothe and Bayfield, 2021)). While this raises questions as to the precise function of Bmc1 catalytic activity on the 5’ end of U6, *in vitro* binding assays showed that catalytic activity of the human MePCE promotes 7SK retention following catalysis (Yang *et al*., 2019). It remains to be found if this extends to other MePCE/Bmc1 targets like U6, and how U6 snRNP assembly may be regulated in species lacking MePCE/Bmc1 and LARP7 homologs.

## Materials and Methods

### Yeast strains and growth

Strains were grown at 32°C in yeast extract with supplements (YES) or Edinburgh Minimal Media (EMM), as indicated. Tag integration and knockouts were generated as described in (Porat *et al*., 2022) (primer sequences provided in supplementary table 1). A list of yeast strains is provided in supplementary table 2.

### Native protein extracts and immunoprecipitation

Native protein extractions and immunoprecipitations were carried out as described in (Porat *et al*., 2022). Protein A-tagged strains were immunoprecipitated with Rabbit IgG-conjugated (MP-Biomedicals, SKU 085594) Dynabeads (Invitrogen, 14301) (Vakiloroayaei *et al*., 2017) and myc- and HA-tagged proteins were immunoprecipitated with Protein A/G beads (GeneBio, 22202B-1) coated with anti-myc antibody (Cell signaling, 2276S) at a dilution of 1:250 or anti-HA antibody (Cell signaling, 3724S) at a dilution of 1:50. Total RNA was isolated from cell extracts with 0.5% SDS, 0.2 mg/mL Proteinase K (Sigma, P2308), 20 mM Tris HCl pH 7.5, and 10 mM EDTA pH 8.0 for 15 minutes at 50°C, followed by phenol: chloroform: isoamyl (25:24:1) extraction and ethanol precipitation. Immunoprecipitated RNA was isolated by incubating beads in 0.1% SDS and 0.2 mg/mL Proteinase K for 30 minutes at 37°C, followed by phenol: chloroform: isoamyl alcohol extraction. For native northern blots, input RNA was extracted in the same manner as immunoprecipitated RNA. Relative immunoprecipitation efficiency was calculated by dividing the IP signal by the input signal. Western blots were performed using anti-myc (Cell signaling, 2276S) at 1:5000, anti-beta actin (Abcam, ab8226) at 1:1250, HRP-conjugated anti-mouse (Cell signaling, 7076) at 1:5000, anti-HA (Cell signaling, 3724S) at 1:1000, HRP-conjugated anti-rabbit (Cell signaling, 7074S) at 1:5000, or HRP-conjugated polyclonal anti-Protein A (Invitrogen, PA1-26853) at 1:5000.

### RNA preparation, northern blotting, 2’-O-methylation detection, and solution hybridization

Total RNA was extracted with hot phenol, separated on 10% TBE-urea polyacrylamide gels, and transferred to positively charged nylon membranes (Perkin Elmer, NEF988001) as per (Porat and Bayfield, 2020). For native RNA extraction to detect U4/U6 duplexes, RNA was extracted with cold phenol, as per (Burke, Butcher and Brow, 2015). Solution hybridization was performed as per (Rodgers *et al*., 2016) and resolved on 9% TBE gels. Probe sequences for ^32^P g-ATP-labeled DNA probes for northern blotting are provided in supplementary table 3. Primer extensions to detect 2’-O-methylation were performed based on protocols from (Zhou *et al*., 2002). Briefly, 5 μg RNA was incubated for 5 minutes at 85°C in a 10 μL reaction containing ^32^P g-ATP-labeled probe, 50 mM Tris HCl pH 7.4, 60 mM NaCl, then transferred to 55°C for 20 minutes to allow the probe to anneal. Reverse transcription was carried out with 1.5 mM (high concentration) or 0.1 mM (limiting concentration) dNTP mix and 2.5 U AMV-RT (NEB, M0277S) and 1 hour incubation at 42°C. cDNA products were separated on 8% TBE-urea sequencing gels, dried, and exposed to Phosphor screens overnight. Relative 2’-O-methylation was calculated by determining the ratio of each RT stop relative to the total signal in each lane (all RT stops and full length U6). 2’-O-methylations were also detected by RNase H (NEB, M0297S) digestion of 2 μg with 25 pmol chimeric RNA-DNA probes, as per (Yu, Shu and Steitz, 1997). Probe sequences targeting C57 and A64 2’-O-methylations are provided in supplementary table 3.

### qRT-PCR and semi-quantitative RT-PCR

1 μg TURBO DNase-treated RNA was reverse transcribed with the iScript cDNA reverse transcription kit (Biorad, 1708890) or 5 U AMV-RT (NEB, M0277S) and gene-specific reverse primers. qRT-PCR was performed with the SensiFAST SYBR No-Rox kit (Bioline, BIO-98005) and 1 μM of each primer, with settings outlined in (Porat *et al*., 2022). For semi-quantitative RT-PCR, cDNA was amplified with Taq polymerase (NEB, MO273L) using the following cycling conditions: 5 min initial denaturation at 94°C, 26 (pud1, alp41) or 27 (rpl1603, bor1) cycles of 30 s at 94°C, 30 s at 50°C, and 1 minute at 72°C, and a final 5 minute extension at 72°C. cDNA was resolved on 10% TBE gels.

### Native yeast extract preparation, native snRNP gels, and glycerol gradient sedimentation

Pellets from 1 L yeast cultures were resuspended to 1 g/mL in AGK400 buffer (10 mM HEPES KOH pH 7.9, 400 mM KCl, 1.5 mM MgCl_2_, 0.5 mM DTT, 1 mM PMSF, and protease inhibitor cocktail (Sigma, P8215)), frozen in liquid nitrogen, and ground to fine powder with a mortar and pestle. Powder was thawed on ice and spun in a JA 25.50 rotor (Beckman) for 16 minutes at 15,000 rpm and the supernatant was subsequently spun in a 70.1 Ti rotor (Beckman) for 45 minutes at 50,000 rpm to pellet ribosomes and heavy molecular weight complexes. Supernatants were flash frozen and stored at −80°C. For native snRNP gels, glycerol with xylene cyanol and bromophenol blue was added to 30 μg cell extract (final glycerol concentration= 10%) and fractionated on 4% 19:1 acrylamide: bis-acrylamide native gels (15 cm x 18 cm) for 220 minutes at 240 V and 4 degrees, then transferred to nylon membranes for northern blotting. For glycerol gradients, cell extracts from 1.0 g frozen cell powder were layered on an 11 mL 10-30% glycerol gradient (50 mM Tris HCl pH 7.4, 25 mM NaCl, 5 mM MgCl_2_) and spun in an SW41Ti rotor (Beckman) for 20 hours at 30,900 rpm. Fractions were collected starting from the top of the gradient and RNA and proteins were extracted with phenol: chloroform: isoamyl alcohol (25:24:1) and TCA precipitation, respectively.

### UV melt curves

UV melt curves were recorded on a Cary BIO 100 spectrometer with a 6 x 6 temperature-controlled cell holder. 2 μL 10 mM U4 and modified or unmodified U6 RNA oligos in 96 μL buffer (10 mM KH_2_PO_4_ pH 7.0 and 200 mM KCl) was heated and cooled from 50°C to 65°C at a rate of 2°C per minute without collecting data, then re-heated and cooled while monitoring absorbance at 260 nm at 1°C intervals. Absorbance at 260 nm at each temperature point was normalized to absorbance at 50°C and absorbance curves were fitted with an equation for one site specific binding with a Hill slope to determine Tm values. RNA sequences are provided in supplementary table 3.

### RNA Seq and intron retention analysis

DNase-treated RNA was rRNA-depleted (Qiagen, 334215) and stranded libraries were prepared by Genome Québec. cDNA libraries were sequenced on a NovaSeq6000 with 150 bp paired-end reads. Reads were aligned to the fission yeast genome (ASM294v2) with Bowtie2 (Langmead and Salzberg, 2012). Intron retention was quantified using IRFinder (version 2.0.1), as per (Lorenzi *et al*., 2021; Schärfen *et al*., 2022). Any introns flagged as having a low sequencing depth or fewer than 4 reads to support splicing were not considered for statistical analysis. Differential intron retention was calculated using DESeq2 (Love, Huber and Anders, 2014).

Sequence extraction for *S. pombe* introns was carried out using BEDTools v2.3.0 (Quinlan and Hall, 2010) and sequences are provided in dataset 1. 5’ splice sites (3 bases in the exon and 6 bases in the intron) and 3’ splice sites (20 bases in the intron and 3 bases in the exon) were scored with MaxEntScan using a maximum entropy model (Yeo and Burge, 2004). Intron free energy of the thermodynamic ensemble (kcal/mol) was calculated using RNAfold v2.5.1 (Lorenz *et al*., 2011).

## Supporting information

Supplementary Dataset S1 Porat et al., 2023

Supplementary Figures Porat et al., 2023

## Data availability

The data supporting the findings of this study are available from the corresponding author upon reasonable request. RNA Seq data have been deposited in NCBI’s Sequence Read Archive (SRA) database under BioProject number PRJNA918556.

## Acknowledgments

We thank Dave Brow for comments on the manuscript. J.P. is supported by a Canada Graduate Scholarship (Doctoral) from the National Sciences and Engineering Research Council of Canada. M.A.B. is supported by a Discovery Grant from NSERC (“Impact of chemical modification of noncoding RNAs on gene expression in *S. pombe”)*. S.D.R. is supported by a Discovery Grant from NSERC (“The molecular mechanism of U6 snRNA activation for pre-mRNA splicing”) and a Sector Innovation Program Grant from GenomeBC (RC18-3517). This research was enabled in part by support provided by the BC DRI Group and the Digital Research Alliance of Canada (alliancecan.ca).

## Author Contributions

J.P. and M.A.B. conceived of the project and designed research, J.P. performed research, J.P. and V.A.S. analyzed the data, J.P. wrote the manuscript, J.P., V.A.S., S.D.R., and M.A.B. edited the manuscript, and M.A.B. and S.D.R. supervised the research and acquired funding.

## References

Abou Assi, H. et al. (2021) ‘2’-O-Methylation can increase the abundance and lifetime of alternative RNA conformational states’, Nucleic Acids Research, 48, pp. 12365–12379. doi: 10.1093/nar/gkaa928.

Awan, A. R., Manfredo, A. and Pleiss, J. A. (2013) ‘Lariat sequencing in a unicellular yeast identifies regulated alternative splicing of exons that are evolutionarily conserved with humans’, Proceedings of the National Academy of Sciences of the United States of America, 110(31), pp. 12762–12767. doi: 10.1073/pnas.1218353110.

Barrass, J. D. et al. (2015) ‘Transcriptome-wide RNA processing kinetics revealed using extremely short 4tU labeling’, Genome Biology, 16 p. 282. doi: 10.1186/s13059-015-0848-1.

Basu, R. et al. (2020) ‘Structure of S. pombe telomerase protein Pof8 C-terminal domain is an xRRM conserved among LARP7 proteins’, RNA Biology, 18(8), pp. 1181–1192. doi: 10.1080/15476286.2020.1836891.

Brow, D. A. and Guthrie, C. (1988) ‘Spliceosomal RNA U6 is remarkably conserved from yeast to mammals’, Nature. Nature Publishing Group, 334(6179), pp. 213–218. doi: 10.1038/334213a0.

Brow, D. A. and Guthrie, C. (1989) ‘Splicing a spliceosomal RNA’, Nature, 337, pp. 14–15. doi: 10.1038/337014a0.

Brown, K. A. et al. (2016) ‘Distinct Dynamic Modes Enable the Engagement of Dissimilar Ligands in a Promiscuous Atypical RNA Recognition Motif’, Biochemistry, 55(51), pp. 7141–7150. doi: 10.1021/acs.biochem.6b00995.

Burke, J. E., Butcher, S. E. and Brow, D. A. (2015) ‘Spliceosome assembly in the absence of stable U4/U6 RNA pairing’, RNA, 21(5), pp. 923–934. doi: 10.1261/rna.048421.114.

Calo, E. et al. (2015) ‘RNA helicase DDX21 coordinates transcription and ribosomal RNA processing’, Nature, 518(7538), pp. 249–253. doi: 10.1038/nature13923.

Chen, W. et al. (2014) ‘Endogenous U2•U5•U6 snRNA complexes in S. pombe are intron lariat spliceosomes’, RNA, 20(3), pp. 308–320. doi: 10.1261/rna.040980.113.

Collopy, L. C. et al. (2018) ‘LARP7 family proteins have conserved function in telomerase assembly’, Nature Communications, 9(1), p. 557. doi: 10.1038/s41467-017-02296-4.

Cosgrove, M. S. et al. (2012) ‘The bin3 RNA methyltransferase targets 7SK RNA to control transcription and translation’, Wiley Interdisciplinary Reviews: RNA. doi: 10.1080/09540121.2017.1344767.

Cvitkovic, I. and Jurica, M. S. (2013) ‘Spliceosome database: a tool for tracking components of the spliceosome’, Nucleic acids research. Nucleic Acids Res, 41(Database issue). doi: 10.1093/NAR/GKS999.

Deragon, J.-M. M. (2020) ‘Distribution, organization an evolutionary history of La and LARPs in eukaryotes’, RNA Biology. Taylor & Francis, 18(2), pp. 1–9. doi: 10.1080/15476286.2020.1739930.

Didychuk, A. L., Butcher, S. E. and Brow, D. A. (2018) ‘The life of U6 small nuclear RNA, from cradle to grave’, RNA. RNA, 24(4), pp. 437–460. doi: 10.1261/rna.065136.117.

Eichhorn, C. D. et al. (2018) ‘Structural basis for recognition of human 7SK long noncoding RNA by the La-related protein Larp7’, Proceedings of the National Academy of Sciences of the United States of America, 115(28), pp. E6457–E6466. doi: 10.1073/pnas.1806276115.

Eysmont, K. et al. (2019) ‘Rearrangements within the U6 snRNA Core during the Transition between the Two Catalytic Steps of Splicing’, Molecular Cell. Cell Press, 75(3), pp. 538–548.e3.

Fica, S. M. et al. (2013) ‘RNA catalyzes nuclear pre-mRNA splicing’, Nature. NIH Public Access, 503(7475), p. 229. doi: 10.1038/NATURE12734.

Frendewey, D. et al. (1990) ‘Schizosaccharomyces U6 genes have a sequence within their introns that matches the B box consensus of tRNA internal promoters’, Nucleic Acids Research, 18(8), pp. 2025–2032. doi: 10.1093/nar/18.8.2025.

Fu, X. et al. (2022) ‘Identification of transient intermediates during spliceosome activation by single molecule fluorescence microscopy’, Proceedings of the National Academy of Sciences of the United States of America. NLM (Medline), 119(48), p. e2206815119. doi: 10.1073/PNAS.2206815119/SUPPL_FILE/PNAS.2206815119.SAPP.PDF.

Ghetti, A., Company, M. and Abelson, J. (1995) ‘Specificity of Prp24 binding to RNA: A role for Prp24 in the dynamic interaction of U4 and U6 snRNAs’, RNA, 1(2), pp. 132–145.

Gildea, M. A., Dwyer, Z. W. and Pleiss, J. A. (2022) ‘Transcript-specific determinants of pre-mRNA splicing revealed through in vivo kinetic analyses of the 1st and 2nd chemical steps’, Molecular cell. Mol Cell, 82(16), pp. 2967–2981.e6. doi: 10.1016/J.MOLCEL.2022.06.020.

Gu, J. et al. (1996) ‘Localization of modified nucleotides in Schizosaccharomyces pombe spliceosomal small nuclear RNAs: Modified nucleotides are clustered in functionally important regions’, RNA, 2, pp. 909–918.

Hasler, D. et al. (2020) ‘The Alazami Syndrome-Associated Protein LARP7 Guides U6 Small Nuclear RNA Modification and Contributes to Splicing Robustness’, Molecular Cell, 77, pp. 1014–1031.e13. doi: 10.1016/j.molcel.2020.01.001.

He, N. et al. (2008) ‘A La-Related Protein Modulates 7SK snRNP Integrity to Suppress P-TEFb-Dependent Transcriptional Elongation and Tumorigenesis’, Molecular Cell, 29(5), pp. 588–599. doi: 10.1016/j.molcel.2008.01.003.

Hu, X. et al. (2020) ‘Quality-Control Mechanism for Telomerase RNA Folding in the Cell’, Cell Reports, 33(13). doi: 10.1016/j.celrep.2020.108568.

Huang, T., Vilardell, J. and Query, C. C. (2002) ‘Pre-spliceosome formation in S.pombe requires a stable complex of SF1-U2AF59-U2AF23’, EMBO Journal, 21(20), pp. 5516–5526. doi: 10.1093/emboj/cdf555.

Huppler, A. et al. (2002) ‘Metal binding and base ionization in the u6 rna intramolecular stemloop structure’, Nature Structural Biology, 9, pp. 431–435. doi: 10.1038/nsb800.

Ishigami, Y. et al. (2021) ‘A single m6A modification in U6 snRNA diversifies exon sequence at the 5’ splice site’, Nature Communications. Nature Publishing Group, 12, p. 3244. doi: 10.1038/s41467-021-23457-6.

Jeronimo, C. et al. (2007) ‘Systematic Analysis of the Protein Interaction Network for the Human Transcription Machinery Reveals the Identity of the 7SK Capping Enzyme’, Molecular Cell, 27, pp. 262–274. doi: 10.1016/j.molcel.2007.06.027.

Ji, C. et al. (2021) ‘Interaction of 7SK with the Smn complex modulates snRNP production’, Nature Communications, 12(1), p. 1278. doi: 10.1038/s41467-021-21529-1.

Jumper, J. et al. (2021) ‘Highly accurate protein structure prediction with AlphaFold’, Nature, 596, pp. 583–589. doi: 10.1038/s41586-021-03819-2.

Kawai, G. et al. (1992) ‘Conformational Rigidity of Specific Pyrimidine Residues in tRNA Arises from Posttranscriptional Modifications That Enhance Steric Interaction between the Base and the 2’-Hydroxyl Group’, Biochemistry, 31(4), pp. 1040–1046. doi: 10.1021/bi00119a012.

Krueger, B. J. et al. (2008) ‘LARP7 is a stable component of the 7SK snRNP while P-TEFb, HEXIM1 and hnRNP A1 are reversibly associated’, Nucleic Acids Research, 36(7), pp. 2219–2229. doi: 10.1093/nar/gkn061.

Kuang, Z., Boeke, J. and Canzar, S. (2017) ‘The dynamic landscape of fission yeast meiosis alternative-splice isoforms’, Genome Research, 27, pp. 145–156.

Kucera, N. J., Hodsdon, M. E. and Wolin, S. L. (2011) ‘An intrinsically disordered C terminus allows the la protein to assist the biogenesis of diverse noncoding RNA precursors’, Proceedings of the National Academy of Sciences of the United States of America, 108(4), pp. 1308–1313. doi: 10.1073/pnas.1017085108.

Langmead, B. and Salzberg, S. L. (2012) ‘Fast gapped-read alignment with Bowtie 2’, Nature Methods, 9(4). doi: 10.1038/nmeth.1923.

Lorenz, R. et al. (2011) ‘ViennaRNA Package 2.0’, Algorithms for Molecular Biology, 6, p. 26. doi: 10.1186/1748-7188-6-26.

Lorenzi, C. et al. (2021) ‘IRFinder-S: a comprehensive suite to discover and explore intron retention’, Genome Biology, 22(1), p. 307. doi: 10.1186/s13059-021-02515-8.

Love, M. I., Huber, W. and Anders, S. (2014) ‘Moderated estimation of fold change and dispersion for RNA-seq data with DESeq2’, Genome Biology, 15(12), p. 550. doi: 10.1186/s13059-014-0550-8.

Lowe, T. M. and Eddy, S. R. (1999) ‘A computational screen for methylation guide snoRNAs in yeast’, Science, 283, pp. 1168–1171.

Markert, A. et al. (2008) ‘The La-related protein LARP7 is a component of the 7SK ribonucleoprotein and affects transcription of cellular and viral polymerase II genes’, EMBO Reports, 9(6), pp. 569–575. doi: 10.1038/embor.2008.72.

Martin-Tumasz, S. et al. (2011) ‘A novel occluded RNA recognition motif in Prp24 unwinds the U6 RNA internal stem loop’, Nucleic Acids Research, 39(17), pp. 7837–7847. doi: 10.1093/nar/gkr455.

Marz, M. et al. (2009) ‘Evolution of 7SK RNA and its protein partners in metazoa’, Molecular Biology and Evolution. Mol Biol Evol, 26(12), pp. 2821–2830. doi: 10.1093/molbev/msp198.

Melangath, G. et al. (2017) ‘Functions for fission yeast splicing factors SpSlu7 and SpPrp18 in alternative splice-site choice and stress-specific regulated splicing’, PLoS ONE, 12, p. e0188159.

Mennie, A. K., Moser, B. A. and Nakamura, T. M. (2018) ‘LARP7-like protein Pof8 regulates telomerase assembly and poly(A)+TERRA expression in fission yeast’, Nature Communications. doi: 10.1038/s41467-018-02874-0.

Montañés, J. et al. (2022) ‘Native RNA sequencing in fission yeast reveals frequent alternative splicing isoforms’, Genome Research, 32, pp. 1215–1227.

Montemayor, E. J. et al. (2018) ‘Architecture of the U6 snRNP reveals specific recognition of 3’-end processed U6 snRNA’, Nature Communications, 9(1). doi: 10.1038/s41467-018-04145-4.

Montemayor, E. J. et al. (2020) ‘Molecular basis for the distinct cellular functions of the Lsm1-7 and Lsm2-8 complexes’, RNA, 26(10), pp. 1400–1413. doi: 10.1261/rna.075879.120.

Naeeni, A. R., Conte, M. R. and Bayfield, M. A. (2012) ‘RNA chaperone activity of human La protein is mediated by variant RNA recognition motif.’, The Journal of biological chemistry, 287(8), pp. 5472–5482. doi: 10.1074/jbc.M111.276071.

Van Nues, R. W. et al. (2011) ‘Box C/D snoRNP catalysed methylation is aided by additional pre-rRNA base-pairing’, EMBO Journal. European Molecular Biology Organization, 30(12), p. 2420. doi: 10.1038/emboj.2011.148.

van Nues, R. W. and Watkins, N. J. (2017) ‘Unusual C’/D’ motifs enable box C/D snoRNPs to modify multiple sites in the same rRNA target region’, Nucleic acids research. Oxford Academic, 45(4), pp. 2016–2028. doi: 10.1093/nar/gkw842.

Páez-Moscoso, D. J. et al. (2018) ‘Pof8 is a La-related protein and a constitutive component of telomerase in fission yeast’, Nature Communications, 9(1), p. 587. doi: 10.1038/s41467-017-02284-8.

Páez-Moscoso, D. J. et al. (2022) ‘A putative cap binding protein and the methyl phosphate capping enzyme Bin3/MePCE function in telomerase biogenesis’, Nature Communications, 13(1), p. 1067. doi: 10.1038/s41467-022-28545-9.

Porat, J. et al. (2022) ‘The methyl phosphate capping enzyme Bmc1/Bin3 is a stable component of the fission yeast telomerase holoenzyme’, Nature Communications, 13(1), p. 1277. doi: 10.1038/S41467-022-28985-3.

Porat, J. and Bayfield, M. A. (2020) ‘Use of tRNA-Mediated Suppression to Assess RNA Chaperone Function’, in Heise, T. (ed.) RNA Chaperones: Methods and Protocols. New York, NY: Springer US, pp. 107–120. doi: 10.1007/978-1-0716-0231-7_6.

Porat, J., Kothe, U. and Bayfield, M. A. (2021) ‘Revisiting tRNA chaperones: New players in an ancient game’, RNA. Cold Spring Harbor Laboratory Press, 27, pp. 543–559. doi: 10.1261/rna.078428.120.

Potashkin, J. and Frendewey, D. (1989) ‘Splicing of the U6 RNA precursor is impaired in fission yeast pre-mRNA splicing mutants’, Nucleic Acids Research, 17(19), pp. 7821–7831. doi: 10.1093/nar/17.19.7821.

Prusiner, P., Yathindra, N. and Sundaralingam, M. (1974) ‘Effect of ribose O(2’)-methylation on the conformation of nucleosides and nucleotides’, BBA Section Nucleic Acids And Protein Synthesis, 366(2), pp. 115–123. doi: 10.1016/0005-2787(74)90325-6.

Quinlan, A. R. and Hall, I. M. (2010) ‘BEDTools: A flexible suite of utilities for comparing genomic features’, Bioinformatics, 26, pp. 841–842. doi: 10.1093/bioinformatics/btq033.

Rader, S. D. and Guthrie, C. (2002) ‘A conserved Lsm-interaction motif in Prp24 required for efficient U4/U6 di-snRNP formation’, RNA, 8, pp. 1378–1392. doi: 10.1017/S1355838202020010.

Reddy, R. and Busch, H. (1988) ‘Small Nuclear RNAs: RNA Sequences, Structure, and Modifications’, in Structure and Function of Major and Minor Small Nuclear Ribonucleoprotein Particles. doi: 10.1007/978-3-642-73020-7_1.

Rinke, J. and Steitz, J. A. (1985) ‘Association of the lupus antigen La with a subset of U6 snRNA molecules’, Nucleic Acids Research, 13(7), pp. 2617–2629. doi: 10.1093/nar/13.7.2617.

Rodgers, M. L. et al. (2016) ‘A multi-step model for facilitated unwinding of the yeast U4/U6 RNA duplex’, Nucleic Acids Research, 44(22), pp. 10912–10928. doi: 10.1093/nar/gkw686.

Schärfen, L. et al. (2022) ‘Identification of Alternative Polyadenylation in Cyanidioschyzon merolae Through Long-Read Sequencing of mRNA’, Frontiers in Genetics, 12, p. 818697.

Shalgi, R. et al. (2014) ‘Widespread Inhibition of Posttranscriptional Splicing Shapes the Cellular Transcriptome following Heat Shock’, Cell Reports, 7, pp. 1362–1370.

Singh, R., Gupta, S., and Reddy, R. (1990) ‘Capping of mammalian U6 small nuclear RNA in vitro is directed by a conserved stem-loop and AUAUAC sequence: conversion of a noncapped RNA into a capped RNA’, Molecular and cellular biology. Mol Cell Biol, 10(3), pp. 939–946. doi: 10.1128/MCB.10.3.939-946.1990.

Singh, M. et al. (2012) ‘Structural Basis for Telomerase RNA Recognition and RNP Assembly by the Holoenzyme La Family Protein p65’, Molecular Cell, 47(1), pp. 16–26. doi: 10.1016/j.molcel.2012.05.018.

Singh, M., Choi, C. P. and Feigon, J. (2013) ‘xRRM: A new class of RRM found in the telomerase La family protein p65’, RNA Biology, 10, pp. 353–359. doi: 10.4161/rna.23608.

Singh, R. and Reddy, R. (1989) ‘γ-Monomethyl phosphate: A cap structure in spliceosomal U6 small nuclear RNA (mRNA splicing/RNA processing/RNA modification)’, Proc. Nadl. Acad. Sci. USA, 86, pp. 8280–8283.

Tani, T. and Ohshima, Y. (1989) ‘The gene for the U6 small nuclear RNA in fission yeast has an intron’, Nature. Nature Publishing Group, 337(6202), pp. 87–90. doi: 10.1038/337087a0.

Tani, T. and Ohshima, Y. (1991) ‘mRNA-type introns in U6 small nuclear RNA genes: Implications for the catalysis in pre-mRNA splicing’, Genes and Development, 5, pp. 1022–1031. doi: 10.1101/gad.5.6.1022.

Vakiloroayaei, A. et al. (2017) ‘The RNA chaperone La promotes pre-tRNA maturation via indiscriminate binding of both native and misfolded targets’, Nucleic acids research, 45(19), pp. 11341–11355. doi: 10.1093/nar/gkx764.

Wang, X. et al. (2020) ‘LARP7-Mediated U6 snRNA Modification Ensures Splicing Fidelity and Spermatogenesis in Mice’, Molecular Cell, 77, pp. 999–1013.e6. doi: 10.1016/j.molcel.2020.01.002.

Warda, A. S. et al. (2017) ‘Human METTL16 is a N 6-methyladenosine (m 6 A) methyltransferase that targets pre-mRNAs and various non-coding RNAs’, EMBO reports, 18, pp. 2004–2014. doi: 10.15252/embr.201744940.

Wilkinson, M. E., Charenton, C. and Nagai, K. (2020) ‘RNA Splicing by the Spliceosome’, Annual Review of Biochemistry, 89, pp. 359–388. doi: 10.1146/annurev-biochem-091719-064225.

Witkin, K. L. and Collins, K. (2004) ‘Holoenzyme proteins required for the physiological assembly and activity of telomerase’, Genes and Development, 18(10), pp. 1107–1118. doi: 10.1101/gad.1201704.

Yang, Y. et al. (2019) ‘Structural basis of 7SK RNA 5’-γ-phosphate methylation and retention by MePCE’, Nature Chemical Biology, 15(2), pp. 132–140. doi: 10.1038/s41589-018-0188-z.

Yeo, G. and Burge, C. B. (2004) ‘Maximum entropy modeling of short sequence motifs with applications to RNA splicing signals’, Journal of Computational Biology, 11, pp. 377–394. doi: 10.1089/1066527041410418.

Yu, Y. T., Shu, M. Di and Steitz, J. A. (1997) ‘A new method for detecting sites of 2’-O-methylation in RNA molecules’, RNA, 3, pp. 324–331.

Zhou, H. et al. (2002) ‘The Schizosaccharomyces pombe mgU6-47 gene is required for 2’-O-methylation of U6 snRNA at A41’, Nucleic Acids Research, 30, pp. 894–902. doi: 10.1093/nar/30.4.894.

