## Supplementary Figures Porat et al., 2023 for "The fission yeast methyl phosphate capping enzyme Bmc1 guides 2’-O-methylation of the U6 snRNA"

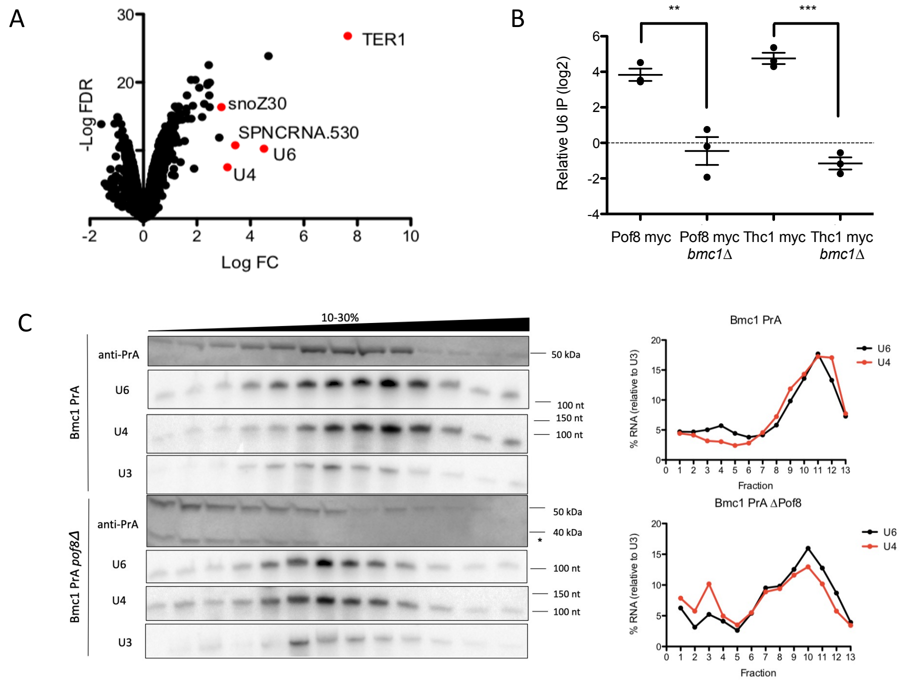


**Figure S1: Bmc1, Pof8, and Thc1 cooperate to bind U6 and U6-associated noncoding RNAs**

A) Enrichment of Bmc1 PrA-associated transcripts compared to an untagged control (n= 3 biological replicates). Axes represent log2 of fold change (FC) and negative log of false discovery rate (FD) (Benjamini-Hochberg adjusted *P* value ≤0.05). Data taken from (Porat *et al.*, 2022).

B) qRT-PCR of U6 in Pof8 myc and Thc1 myc immunoprecipitates, normalized to immunoprecipitation from an untagged strain (mean± standard error, two-tailed unpaired *t* test, $^{**}p<0.01$, $^{***}p<0.001$) (n= 3 biological replicates).

C) Glycerol gradient sedimentation of PrA-tagged Bmc1, U4, U6, and U3 from wild type (Bmc1 PrA) and *pof8Δ* strains. Cleavage products are indicated with an asterisk. U4 and U6 signals were normalized to U3 for calculating relative migration in the gradient.

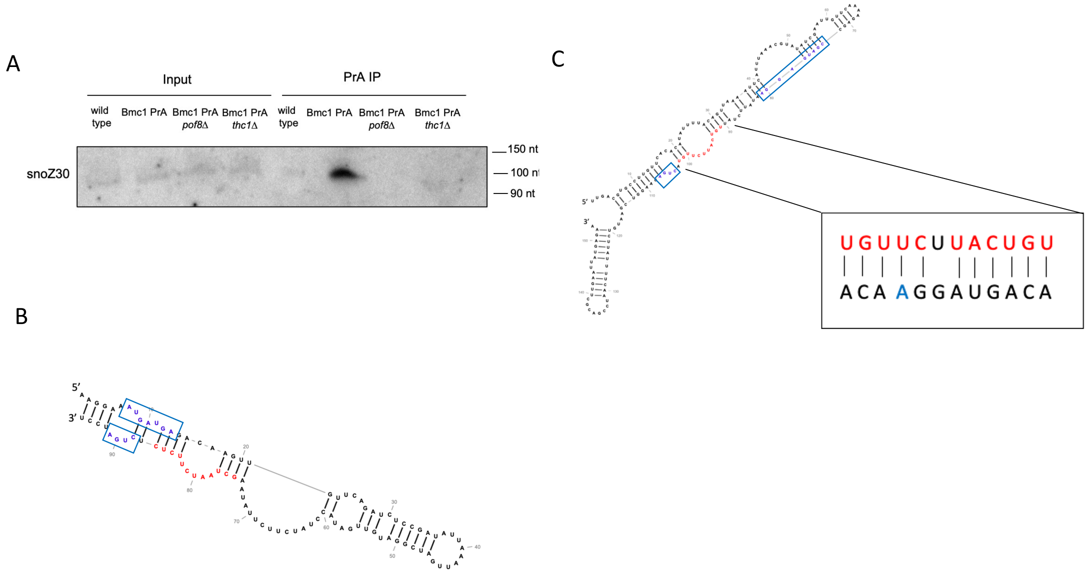


**Figure S2: snoZ30 and sno530 are Bmc1-interacting, U6-modifying snoRNAs**

A) Northern blot analysis of snoZ30 in total RNA and PrA immunoprecipitates from an untagged strain (wild type) and wild type and knockout PrA-tagged strains.

B) Secondary structure prediction (Gruber *et al.*, 2008) of snoZ30. C and D boxes are indicated in blue and U6-binding site is indicated in red.

C) Secondary structure prediction (Gruber *et al.*, 2008) of sno530. C and D boxes are indicated in blue and U6-binding site is indicated in red. Inset: U6-interacting region, highlighting Watson Crick and non-Watson Crick base pairs with U6 (black). A64 is indicated in blue.

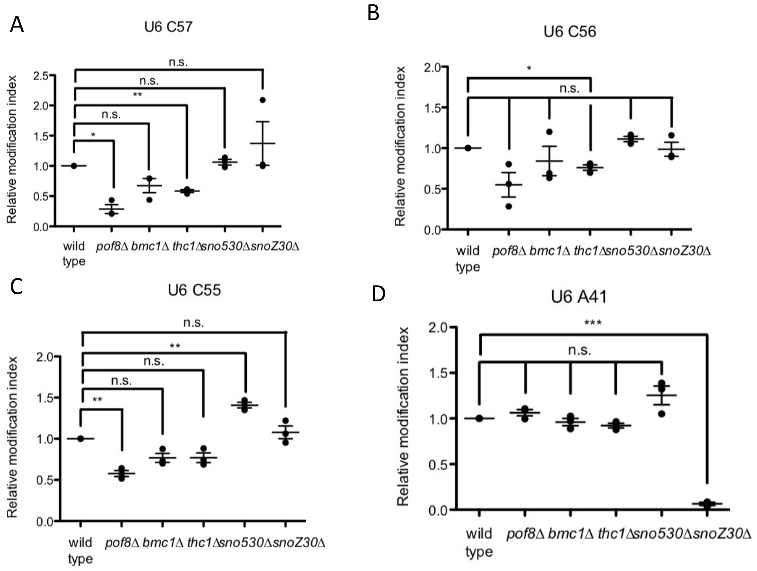


**Figure S3: Bmc1, Pof8, and Thc1 influence 2'-O-methylation of U6**
Quantification of relative 2'-O-methylation-induced reverse transcriptase stops, compared to a wild type strain, for C57 (A), C56 (B), C55 (C),and A41 (D) (mean± standard error, two-tailed paired *t* test, **p*<0.05, ***p*<0.01, *** *p*<0.001) (n= 3 biological replicates).

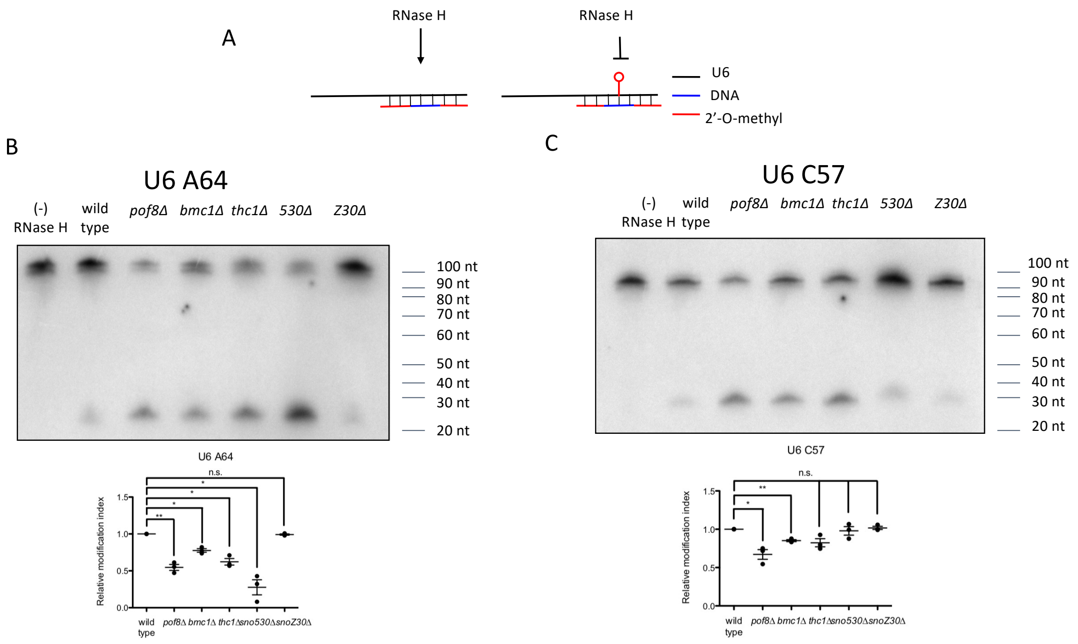


**Figure S4: RNase H cleavage validates 2'-O-methylation of U6 at A64 and C57**
A) Schematic of RNase H cleavage assay to detect 2'-O-methylations (Yu, Shu and Steitz, 1997; Calo *et al.*, 2015).
B-C) Northern blot analysis and quantification of 2'-O-methylation at A64 (B) and C57 (C). Relative modification is expressed as a fraction of the cleaved band relative to total U6 (mean± standard error, two-tailed paired *t* test, **p*<0.05, ***p*<0.01) (n= 3 biological replicates).

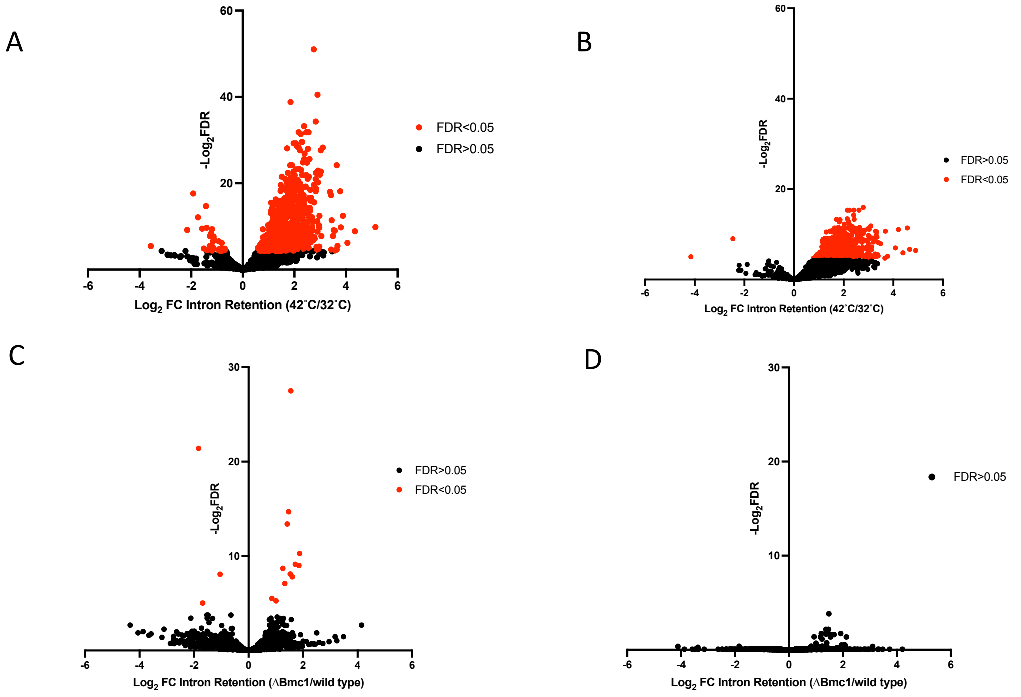


**Figure S5: Heat shock and Bmc1 deletion lead to changes in intron retention.**
A-B) Changes in intron retention in wild type (A) and *bmc1Δ* (B) strains grown at 32˚C or heat shocked for 15 minutes at 42˚C (n=3 biological replicates). Axes represent log2 of fold change (FC) and negative log2 of false discovery rate (FD) (Benjamini-Hochberg adjusted *P* value ≤0.05). C-D) Changes in intron retention in wild type and *bmc1Δ* strains grown at 32˚C (A) or heat shocked for 15 minutes at 42˚C (B) (n=3 biological replicates). Axes represent log2 of fold change (FC) and negative log2 of false discovery rate (FDR) (Benjamini-Hochberg adjusted *P* value ≤0.05).

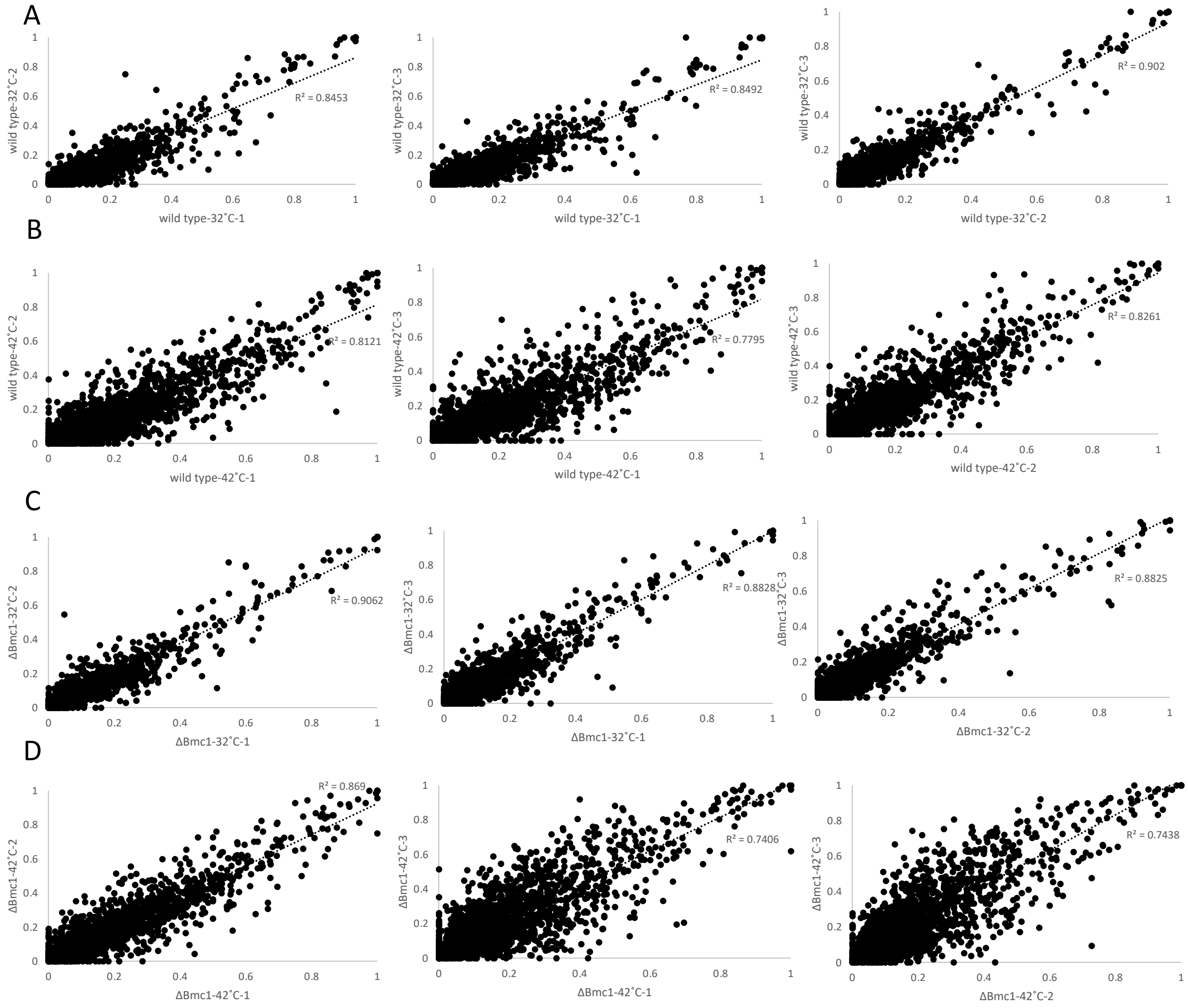


**Figure S6: Correlations between replicates for RNA-Seq of wild-type and *bmc1Δ* strains with and without heat shock.**

Intron retention values for wild type RNA Seq samples grown at 32˚C (A) and 42˚C (B) and ΔBmc1 cells grown at 32˚C (C) and 42˚C (D). $R^2$ values are displayed.

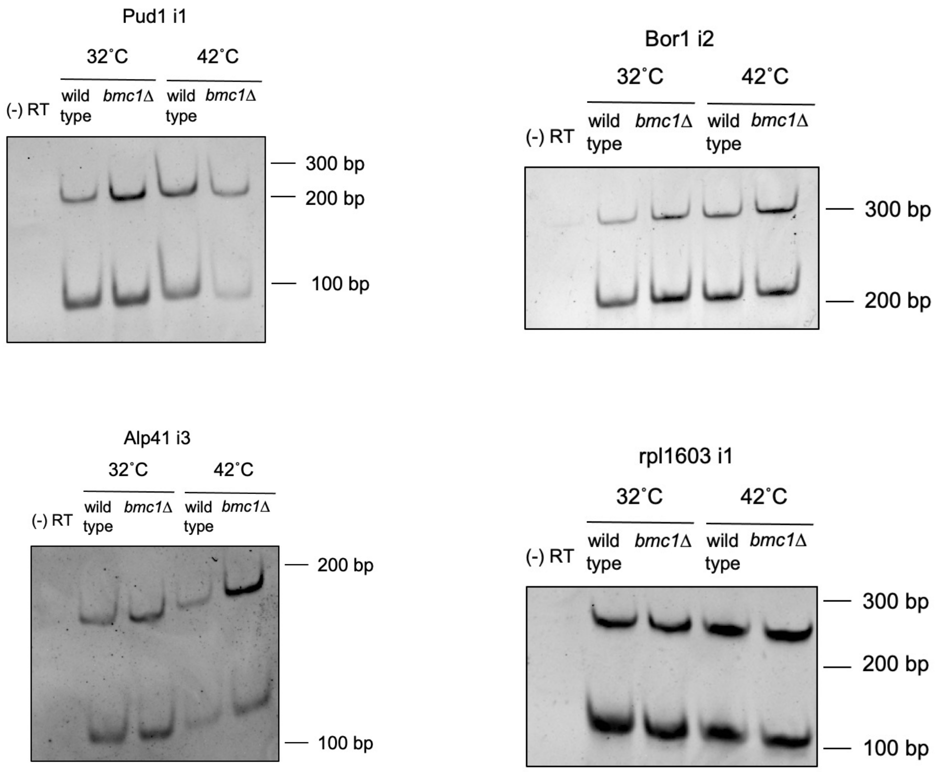


**Figure S7: Semi-quantitative RT-PCR validation of heat shock- and Bmc1-sensitive intron retention events.**

Representative gels contributing to quantifications in figure 5B.

**Table S1: Primer sequences for the creation of tagged and knockout *S. pombe* strains**

| Primer | Sequence |
|---|---|
| 5' Thc1 myc SalI For | 5' GCGCGTCGACATATTCATTGAAGTTGTAGTTTTAGCTTATTTGGTACCA AAAG 3' |
| 5' Thc1 myc BamHI Rev | 5' GCGCGGATCCGACCTGAAGTCAAGAAATTTTGTAGTGGAATAATTCTT TTC 3' |
| 3' Thc1 myc SacI For | 5'GCGCGAGCTCACAGATAAATTAGAACACAGCTTAAACTTACCGGAAAAA TTAATC 3' |
| 3' Thc1 myc SacII Rev | 5' GCGCCCGCGGTTAGATAGCCATGAATAAATAATATCAAGAATATAATC ATCTAAGC 3' |
| 5' Prp24 myc XhoI For | 5' CGCGCTCGAGCCTTAGCTAAAAGCTTTGAGACTACTGAGTCAAATAAA ATG 3' |
| 5' Prp24 myc BamHI Rev | 5' GCGCGGATCCGTTTTAAAAACATTTTCCTAAAATCATCGTTGCTTTTAG GTGCATC 3' |
| 3' Prp24 myc SacI Rev | 5' GCGCGAGCTCTCAGCAATAATTAGATTGAATGATTAAAAAAATATTGAA AACC 3' |
| 3' Prp24 myc SacII Rev | 5' CGCGCCGCGGTGAATATCATTAAAACTCCTTCATTTACTCTTGATGAAA AAATTCAAGAGTC 3' |
| 5' Thc1 KO SalI For | 5' GCGCGTCGACATAATAACTTTGCTTACGATTAATAGACAAATGAATGC TG 3' |
| 5' Thc1 KO BglII Rev | 5' GCGCAGATCTTCTTCAAAACTTTTGGTACCAAATAAGCTAAAACTACA A 3' |
| 3' Thc1 KO ClaI For | 5' GCGCATCGATACAGATAAATTAGAACACAGCTTAAACTTACCGGAAAA A 3' |
| 3' Thc1 KO SacII Rev | 5' GCGCCCGCGGCTTTACTTAAACAGAGAAAAAAAAAACATCTGGAGA C 3' |
| 5' sno530 KO SalI For | 5' GCGCGTCGACGCATGTAAACTGTTTCACGACTTAACGGATCATATG G 3' |
| 5' sno530 KO BglII Rev | 5' GCGCAGATCTATTGACAACAATCAACTGCGTGTTTTATTTACTTTTAAG A 3' |
| 3' sno530 KO ClaI For | 5' GCGCATCGATTCATACAAAAAACTGAGGCTATTTGCTTTAACTGTAGC T 3' |
| 3' sno530 KO SacII Rev | 5' GCGCCCGCGGTGAACCTCAAGTCCCGCAAAATACTTCCAATAATAATT TG 3' |
| 5' snoZ30 KO BamHI For | 5' GCGCGGATCCATTTCTTTGCATCCTTTAATATACTTTCCAATTCATTAA G 3' |
| 5' snoZ30 KO AscI Rev | 5' GGCGCGCGCGCCCTCCTGGTCGACTTTATGGAACTAAAGGGTTACAA TTG 3' |
| 3' snoZ30 KO SacI For | 5' GCGCGAGCTCATCAAAGTAACTTTCTCGGAGAAAGGTCGAGCAATTG TAAT 3' |
| 3' snoZ30 KO ClaI Rev | 5' GCGCATCGATACAAATAAATACGATTAGTCTTAAGTTAATTTAGCACAA AGATTTAAAGTC 3' |

**Table S2: List of yeast strains used in this study**

| Figure | Strain | Description | Full genotype | Source |
|---|---|---|---|---|
| 1a, 1c, 2, 3, S1a, S1b, S2a, S3, S4, S5 | y12088 | wild type | *h+ ura4-D18 leu1-32* | Lab stock |
| 1, 2c, S1a, S2a | yJP001 | *bmc1-PrA* | *h- ura4-D18 bmc1⁺::bmc1-PrA- kanMX6* | Porat et al., 2022 |
| 1a, 1b, S2a | yJP011 | *bmc1-PrA pof8Δ* | *h- leu1-32 ura4-D18 his3-D1 bmc1⁺::bmc1-PrA-kanMX6 pof8Δ::natMX6* | Porat et al., 2022 |
| 1a, S2a | yJP027 | *bmc1-PrA thc1Δ* | *his3-D1 bmc1⁺::bmc1-PrA-kanMX6 thc1Δ::bleMX6* | This study |
| 1c, 2a, 2d, 5, S3, S4 | TN12118a | *pof8Δ* | *h- leu1-32 ura4-D18 his3-D1 pof8Δ::natMX6* | Mennie et al., 2018 and Porat et al., 2022 |
| 1c, 2a, 2d, 3, 4, S3, S4 | yJP022 | *bmc1Δ* | *h+ ura4-D18 leu1-32 bmc1Δ::bleMX6* | Porat et al., 2022 |
| 1c, 2a, 2d, S3, S4 | yJP026 | *thc1Δ* | *h+ ura4-D18 thc1Δ:: bleMX6* | This study |
| 1c, 2a, 2d, S3, S4 | yJP030 | *sno530Δ* | *h+ ura4-D18 sno530Δ::bleMX6* | This study |
| 1c, S3, S4 | yJP031 | *snoZ30Δ* | *h- ura4-D18 snoZ30Δ::bleMX6* | This study |
| S2b, S2c | AM16932 | *pof8-myc* | *h- leu1-32 ura4-D18 his3-D1 pof8⁺::13myc-kanMX6* | Mennie et al., 2018 |
| S2b, S2c | yJP023 | *pof8-myc bmc1Δ* | *leu1-32 ura4-D18 his3-D1 pof8⁺::13myc-kanMX6 bmc1Δ::bleMX6* | Porat et al., 2022 |
| S2b, S2c | yJP029 | *thc1-myc* | *h- leu1-32 ura4-D18 his3-D1 thc1⁺::13myc-kanMX6* | This study |
| S2b, S2c | yJP046 | *thc1-myc bmc1Δ* | *leu1-32 ura4-D18 his3-D1 thc1⁺::13myc-kanMX6 bmc1Δ::bleMX6* | This study |
| 2g | yJP048 | *prp24-myc* | *h- leu1-32 ura4-D18 his3-D1 prp24⁺::13myc-kanMX6* | This study |

| | | | | |
|---|---|---|---|---|
| 2g | yJP049 | *prp24-myc pof8Δ* | *leu1-32 ura4-D18 his3-D1 prp24⁺::13myc-kanMX6 pof8Δ::natMX6* | This study |
| 2g | yJP050 | *prp24-myc bmc1Δ* | *leu1-32 ura4-D18 his3-D1 prp24⁺::13myc-kanMX6 bmc1Δ::bleMX6* | This study |

**Table S3: List of primer and RNA sequences used in this study**

| Figure | Probe | Sequence |
|---|---|---|
| 1a, S1b | U6 RT-PCR For | 5' CGGATCACTTTGGTCAAATTG 3' |
| | U6 RT-PCR Rev | 5' CTCTCAATGTCGCAGTGTCATC 3' |
| 1a | 530 RT-PCR For | 5' ATGAGGAATATTCTATTGTCATTC 3' |
| | 530 RT-PCR Rev | 5' AACAATTCGATATACGTTTAAATG 3' |
| 1b, 1c, 2a, 2c, 2d, 2g, 4b, 4e, 5b, 5d, S2c, S2d | U6 northern, primer extension, solution hybridization | 5' AATGGGTTTTCTCTCAATGTCGCAG 3' |
| 1b, 2a, 2d, 2g | U4 northern and solution hybridization | 5' GTTGGAGCGGTCAGGGTAATAGT 3' |
| S2a | snoZ30 northern | 5' GGAGATCTGAACAACTTGTCTCATC 3' |
| 2a | U1 northern | 5' GCTGCAGAAACTCATGCCAGGTAAGT 3' |
| 2a | U2 northern | 5' TGCCAGTAGTGCAATAGCAAGAACAC 3' |
| 2a | U3 northern | 5' ACACGTCAGAAAACACCAGCTGCCC 3' |
| 2a | U5 northern | 5' GATTACAAAAACTATACAGTCAAATTAGCAC 3' |
| 2f | Unmodified U6 oligo | 5' rUrGrGrCrCrCrCrUrGrCrArCrArArGrGrArUrGrArCrA 3' |
| 2f | A64m U6 oligo | 5' rUrGrGrCrCrCrCrUrGrCrArCrAmArGrGrArUrGrArCrA 3' |
| 2f | U4 oligo | 5' rArUrCrUrUrUrGrUrGrCrArCrGrGrGrUrArU 3' |
| S4b | U6 A64 RNase H chimeric oligo | 5' mUmCCTTGmUmGmCmAmGmGmGmGmCmCmAmU 3' |
| S4c | U6 C57 RNase H chimeric oligo | 5' mGmCAGGGmGmCmCmAmUmGmCmUmAmAmUmC 3' |

**Dataset S1 (separate file).** Intron retention (IR) ratio values and intron features for wild type (y12088) and ΔBmc1 RNA Seq analysis at 32˚C and 42 ˚C. Introns with more than 4 reads supporting splicing in all biological replicates are included.

**Supplemental References**

Calo, E. *et al.* (2015) 'RNA helicase DDX21 coordinates transcription and ribosomal RNA processing', *Nature*, 518(7538), pp. 249–253. doi: 10.1038/nature13923.

Gruber, A. R. *et al.* (2008) 'The Vienna RNA Websuite', *Nucleic Acids Research*, 36(suppl_2), pp. W70–W74. doi: 10.1093/nar/gkn188.

Porat, J. *et al.* (2022) 'The methyl phosphate capping enzyme Bmc1/Bin3 is a stable component of the fission yeast telomerase holoenzyme', *Nature Communications*, 13(1), p. 1277. doi: 10.1038/S41467-022-28985-3.

Yu, Y. T., Shu, M. Di and Steitz, J. A. (1997) 'A new method for detecting sites of 2′-O-methylation in RNA molecules', *RNA*, 3, pp. 324–331.
